# Repurposing chloramphenicol acetyltransferase for a robust and efficient designer ester biosynthesis platform

**DOI:** 10.1101/2020.11.04.368696

**Authors:** Hyeongmin Seo, Jong-Won Lee, Richard J. Giannone, Noah J. Dunlap, Cong T. Trinh

## Abstract

Robust and efficient enzymes are essential modules for metabolic engineering and synthetic biology strategies across biological systems to engineer whole-cell biocatalysts. By condensing an acyl-CoA and an alcohol, alcohol acyltransferases (AATs) can serve as an interchangeable metabolic module for microbial biosynthesis of a diverse class of ester molecules with broad applications as flavors, fragrances, solvents, and drop-in biofuels. However, the current lack of robust and efficient AATs significantly limits their compatibility with heterologous precursor pathways and microbial hosts. Through bioprospecting and rational protein engineering, we identified and repurposed chloramphenicol acetyltransferases (CATs) from mesophilic prokaryotes to function as robust and efficient AATs compatible with at least 21 alcohol and 8 acyl-CoA substrates for microbial biosynthesis of linear, branched, saturated, unsaturated and/or aromatic esters. By plugging the best engineered CAT (CATec3 Y20F) into the gram-negative mesophilic bacterium *Escherichia coli*, we demonstrated that the recombinant strain could effectively convert various alcohols into desirable esters, for instance, achieving a titer of 13.9 g/L isoamyl acetate with 95% conversion by fed-batch fermentation. The recombinant *E. coli* was also capable of simulating the ester profile of roses with high conversion (> 97%) and titer (> 1 g/L) from fermentable sugars at 37°C. Likewise, a recombinant gram-positive, cellulolytic, thermophilic bacterium *Clostridium thermocellum* harboring CATec3 Y20F could produce many of these esters from recalcitrant cellulosic biomass at elevated temperatures (>50°C) due to the engineered enzyme’s remarkable thermostability. Overall, the engineered CATs can serve as a robust and efficient platform for designer ester biosynthesis from renewable and sustainable feedstocks.

## Introduction

Metabolic engineering and synthetic biology approaches enable controllable manipulation of whole-cell biocatalysts to produce valuable chemicals from renewable and sustainable feedstocks and organic wastes in a rapid and efficient manner, helping reduce our reliance on the conventional petroleum-based chemical synthesis (1–3). Enzymes are essential modules that facilitate interchangeable design of metabolic pathways to perform desirable functions across microbial hosts (4, 5). Critical to the optimal design of heterologous pathways is the availability of robust and efficient enzymes that are compatible with various endogenous pathways and microbial hosts to enable combinatorial biosynthesis of desirable molecules.

One important example requiring broad enzymatic compatibility, robustness, and efficiency is engineering of microbial systems to produce a diverse class of esters. Industrially, esters have versatile applications as flavors (6, 7), fragrances (8), solvents (9), and fuels (10–13). In nature, volatile esters are formulated by an alcohol acyltransferase (AAT, EC 2.3.1.84) that condenses an alcohol and an acyl-CoA in a thermodynamically favorable reaction, providing flavors and fragrances in ripening fruits (14) and fermenting yeasts (7) and having an ecological role in pollination (15). Inspired by nature, most of the metabolic engineering and synthetic biology strategies have deployed the eukaryotic AATs originating from plants or yeasts for microbial biosynthesis of target esters (8, 9, 16, 17). However, these eukaryotic AATs are lack of robustness, efficiency, and compatibility as they commonly exhibit poor enzyme expression (18), solubility (19), and thermostability (20) in microbes, thus limiting optimal microbial production of esters. In addition, limited knowledge on substrate profiles and specificities of AATs often requires laborious bioprospecting of AATs for individual target esters (8, 9, 13, 17, 18). Therefore, developing robust and efficient AATs compatible with multiple pathways and microbial hosts is critical to expand biological routes for designer ester biosynthesis.

Chloramphenicol O-acetyltransferase (CAT, EC 2.3.1.28) is an antibiotic resistance enzyme that detoxifies chloramphenicol and derivative antibiotics, which inhibit protein elongation in organisms and cause cell death, by acetylation (21). Organisms resist to this potent drug by harboring CATs that display nearly perfect catalytic efficiency at recruiting an acetyl-CoA(s) to detoxify chloramphenicol (22, 23). In nature, the *cat* gene is one of the most widespread genetic elements (24), expressing a functional enzyme in a wide range of organisms including plants (25), animals (26), and bacteria (27). Interestingly, when being used as antibiotic selection in a recombinant *Escherichia coli*, some CATs exhibited substrate promiscuity resulting in unexpected production of esters (8, 28). Recently, an engineered cellulolytic thermophile *Clostridium thermocellum (Hungateiclostridium thermocellum)* harboring a CAT derived from a mesophile *Staphylococcus aureus* (CATsa) was capable of producing isobutyl acetate from cellulose at elevated temperatures (29, 30). Here, we investigated whether novel CATs can be identified and repurposed to function as a robust and efficient AAT compatible with a wide range of pathways and microbial hosts to enable designer ester biosynthesis.

## Results and discussion

### Identification of beneficial mutations for repurposing CATs as robust and efficient AATs

We first examined whether CATs could be engineered for enhanced robustness and efficiency towards smaller chain alcohols different from the native substrate chloramphenicol (Fig. 1A). To guide CAT engineering, we created a protein homology model of a thermostable and promiscuous CATsa of *S. aureus* from the available protein structures (29, 31–33) and analyzed its stability and binding affinity with smaller chain alcohols and acetyl-CoA required for ester biosynthesis (Figs. 1B, 1C). As a representative alcohol, isobutanol was chosen for docking simulation and experimental characterization of CAT variants for robustness and efficiency because this alcohol is much smaller and very different from chloramphenicol and can be efficiently produced by organisms (34, 35). Affinity and stability screening via *in silico* mutagenesis of selected CATsa binding pocket residues suggested some candidates potentially altering the isobutanol binding (Fig. S1). By experimentally constructing and characterizing the top 26 CATsa variants using an *in vivo* microbial screening assay (36, 37), we identified the CATsa Y20F variant to exhibit the most significant increase in conversion of isobutanol to isobutyl acetate, about 43-fold higher than its wild type (Table S1).

**Figure 1.**
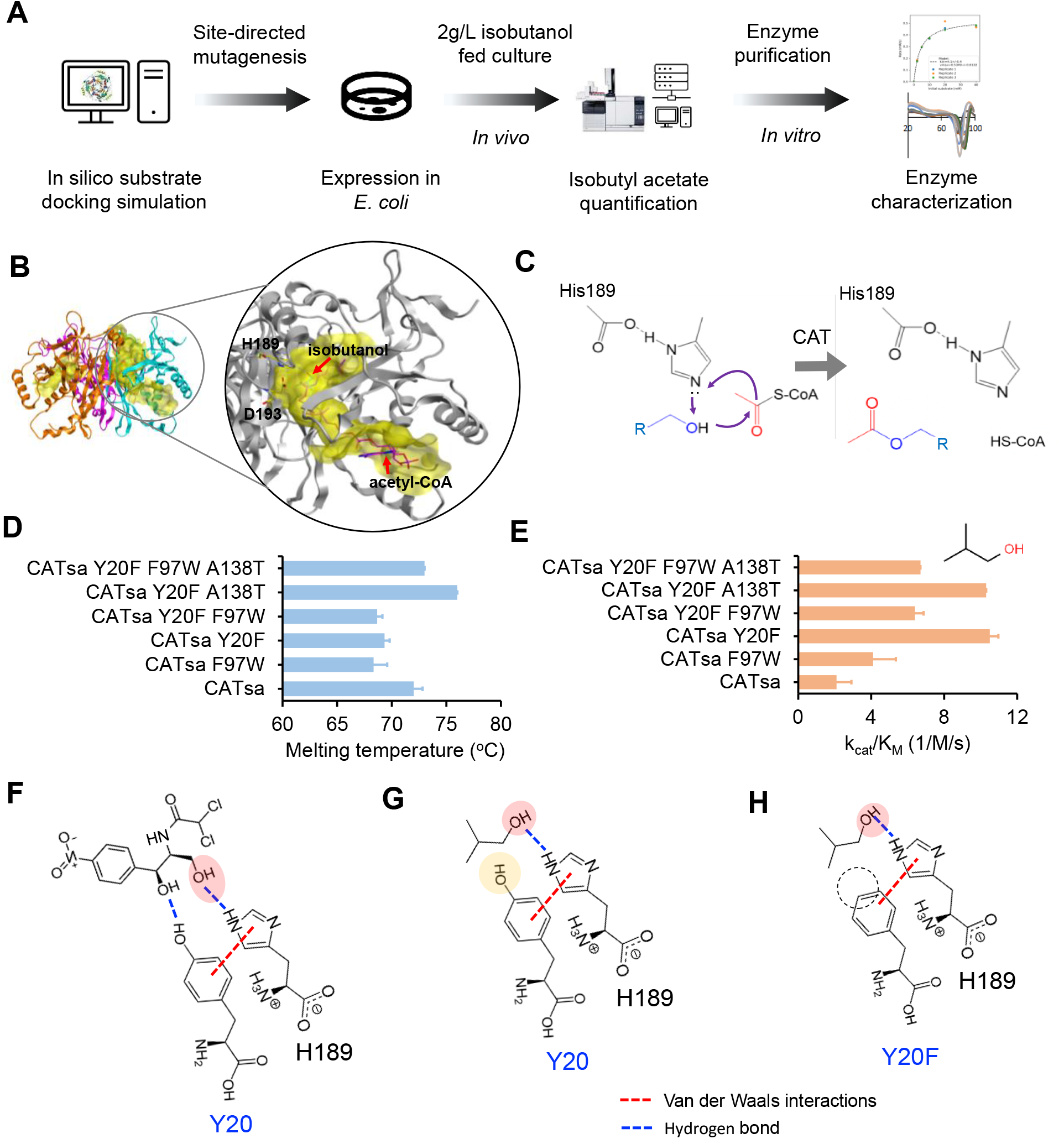
Protein engineering of CAT derived from *Staphylococcus aureus* (CATsa) to improve efficiency and robustness towards isobutanol. **(A)** A schematic workflow of CATsa protein engineering. **(B)** Protein homology model of CATsa. **(C)** Proposed reaction mechanism of ester biosynthesis from an alcohol and acetyl-CoA. **(D)** Melting temperatures of CATsa and its variants. **(E)** Catalytic efficiencies of CATsa and its variants towards isobutanol. In panels **D-E**, each value represents mean ± 1 stdev from at least three biological replicates. For each *in vitro* assay, 2 mM acetyl-CoA was used as a co-substrate. **(F-H)** Interaction of the amino acid residue 20 (i.e., Y20 or Y20F), His-189, and alcohol (i.e., chloramphenicol or isobutanol) in a transition state. In panel **F**, the active site His-189 interacts with the native residue Y20 and chloramphenicol. In panel **G**, the native substrate chloramphenicol in panel **F** is swapped with isobutanol. In panel **H**, Y20 in panel **G** is replaced with Y20F.

We next characterized the robustness and efficiency of CATsa Y20F *in vitro*. Since previous studies demonstrated that CATsa F97W improved the activity towards isobutanol (29) and CATsa A138T increased thermostability (38), we also investigated whether the combinatorial mutagenesis from these single beneficial CATsa variants enhanced the enzyme performance (Figs. 1D, 1E). We found that CATsa Y20F improved the catalytic efficiency over the wildtype CATsa and CATsa F97W by 5.0- and 2.5-fold, respectively (Fig. 1E), while the melting temperature slightly decreased from 71.2 to 69.3°C (Fig. 1D). Among the combinatorial mutagenesis, CATsa Y20F A138T exhibited both the highest melting temperature (76 ± 0.0°C) and catalytic efficiency towards isobutanol (10.3 ± 1.2, 1/M/s), respectively. It has been suggested that the hydrogen bonds between chloramphenicol, the catalytic site His-189, and Tyr-20 at a transition state are critical for high catalytic efficiency of the CAT, specifically by constraining the binding of chloramphenicol in the correct orientation(39) (Fig. 1F). Since the Y20F mutation retains the aromatic ring that contributes to tautomeric stabilization of His-189, the catalytic imidazole can still interact with the smaller chain alcohols flexibly (Figs. 1G, 1H), likely contributing to the enhanced enzymatic activity observed for isobutyl acetate biosynthesis (Fig. 1E).

### Bioprospecting of CATs

To explore whether the high thermostability and alcohol promiscuity of CATs occurred in nature, we used bioinformatic analysis to select a library of 27 CATs representing type-A and type-B that are structurally distinctive (24) and derived from different organisms (Fig. 2A, Table S2). We synthesized these CATs, expressed, purified, and characterized their melting temperatures and promiscuous activities towards isobutanol (Figs. 2A, S2). Most of the CATs showed melting temperatures higher than 60°C except CAT_GEO (T_m_ = 43.5°C) (Table S2). Among these CATs, CAT1_ECOLIX (CATec1), CAT3_ECOLIX (CATec3), CATsa, CAT_KLEPS (CATkl), CAT2_ECOLIX (CATec2), CAT_HAEIF (CATha), and CAT_CLOBU (CATcb) exhibited the highest specific activities towards isobutanol at 50°C (Fig. S2B). Five out of these seven most isobutanol-active CATs were evolutionarily related, suggesting that their activities towards isobutanol might be influenced by their unique structural features. We further evaluated kinetic thermostability of these seven most isobutanol-active CATs by measuring their activity losses after one-hour incubation at elevated temperatures of 50, 55, 60, 65, and 70°C (Fig. S3). Remarkably, CATec3 and CATec2, derived from a mesophilic *E. coli*, could retain more than 95% of the activity at 70°C while the residual activity of CATsa rapidly decreased. Structural analysis suggested that CATec3 has stronger interaction energy than CATsa (Fig. S4).

**Figure 2.**
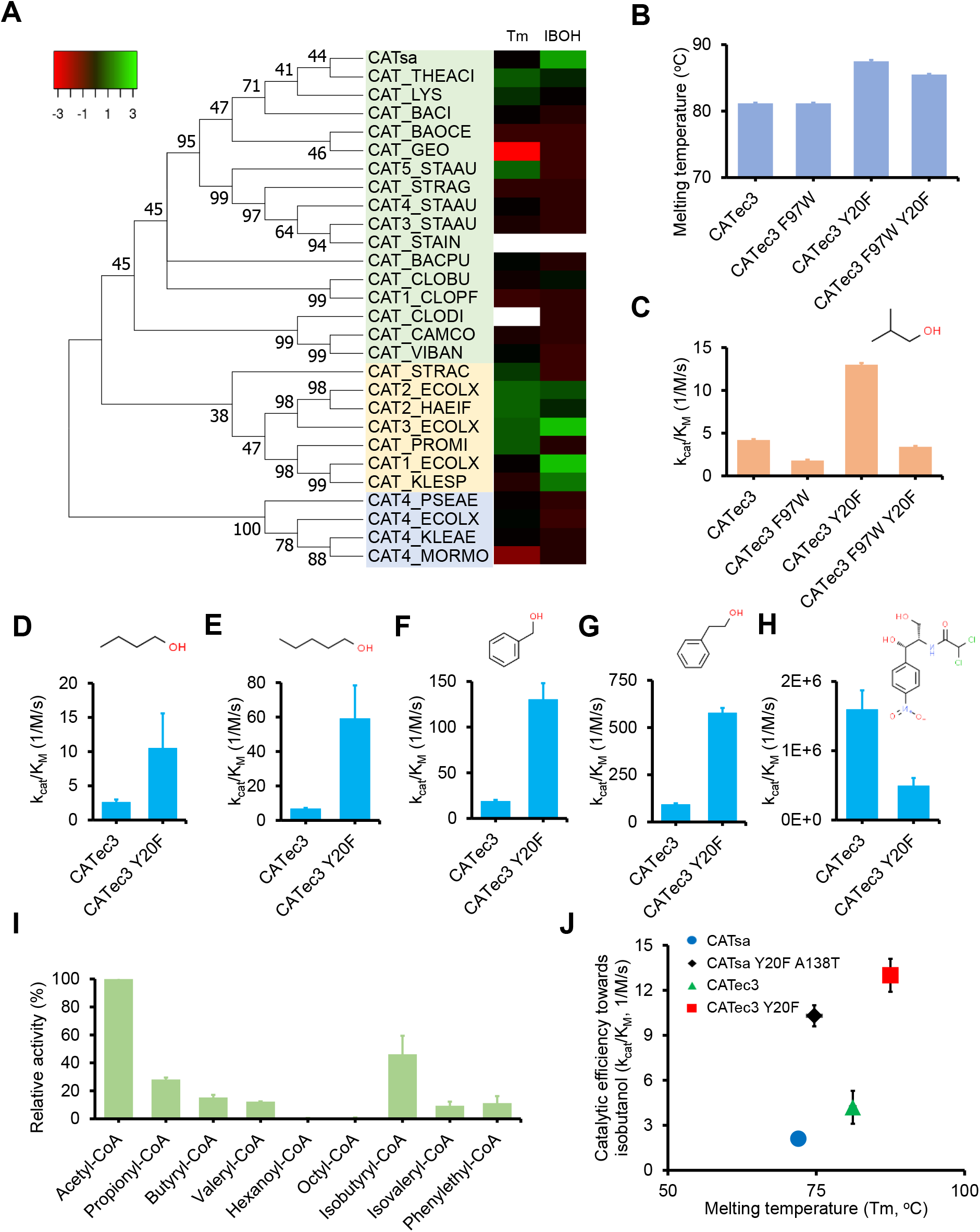
Bioprospecting and protein engineering of CATs. **(A)** A phylogenetic tree of 28 CATs and heat map of their melting temperatures (Tm) and activities towards isobutanol (IBOH). The numbers in the phylogenetic tree represent bootstrapping values (%) from 1000 bootstrap replicates. **(B)** Melting temperatures of CAT3_ECOLIX (CATec3) variants. **(C)** Catalytic efficiencies of CATec3 and its variants towards isobutanol. **(D-H)** Comparison of catalytic efficiencies of CATec3 and CATec3 Y20F towards **(D)** butanol, **(E)** pentanol, **(F)** benzyl alcohol, **(G)** 2-phenylethyl alcohol, and **(H)** chloramphenicol. In panels **B-H**, each value represents mean ± 1 stdev from at least three biological replicates. **(I)** Relative activity of CATec3 Y20F towards acyl-CoAs as compared to acetyl-CoA. Isobutanol (400mM) was supplemented as the co-substrate. Each value represents mean ± 1 stdev from six biological replicates. **(J)** Improvements on melting temperatures and catalytic efficiencies of engineered CATs towards isobutanol. Each value represents mean ± 1 stdev from at least three biological replicates.

### Engineering CAT for enhanced robustness, efficiency, and compatibility

Discovery of high thermostability and promiscuous activity of CATec3 towards a much smaller alcohol (i.e., isobutanol) (Figs. S2, S3) prompted us to investigate whether the beneficial mutations discovered in CATsa could be introduced into CATec3 to further improve its performance. We constructed CATec3 variants harboring single and combinatorial mutations of F97W and Y20F. Since the homolog residue of Ala-138 does not exist in CATec3, its mutation was not constructed and characterized. Interestingly, the results showed that the CATec3 Y20F variant improved not only the catalytic efficiency (13.0 ± 0.2, 1/M/s), about 3.3-fold higher than its wildtype, but also the melting temperature (87.5 ± 0.5°C). Among all the CATs characterized, CATec3 Y20F is the most thermostable and isobutanol-active (Table S3).

Motivated by the flexible structure introduced by the Y20F mutation towards various alcohols (Figs. 1H, S5), we further examined whether CATec3 Y20F exhibited enhanced efficiency and compatibility towards a library of 16 linear, branched, saturated, unsaturated, and aromatic alcohols with acetyl-CoA as a co-substrate (Table 1). We found that CATec3 Y20F was more efficient towards bulky and long-chain alcohols that are more hydrophobic than short-chain alcohols (Table 1), likely due to a stronger binding affinity (Fig. S5D). As compared to the wild type CATec3 against a representative set of six alcohols that can be naturally synthesized by organisms (Fig. S6), we found that the CATec3 Y20F variant exhibited much higher catalytic efficiency towards butanol by 4.0-fold (Fig. 2D), pentanol by 8.8-fold (Fig. 2E), benzyl alcohol by 6.9-fold (Fig. 2F), and phenylethyl alcohol by 6.2-fold (Fig. 2G), respectively. In contrast, the catalytic activity of CATec3 Y20F towards the native substrate chloramphenicol decreased about 3.2-fold as compared to the wild type activity (Fig. 2H).

**Table 1.** Catalytic efficiency of CATec3 Y20F towards multiple alcohol substrates. The catalytic efficiency was measured from the kinetic reactions performed at 50°C. The co-substrate, acetyl-CoA, was supplemented at the saturated concentration of 2 mM. The values represent average ± standard deviation from at least three biological replicates. *Calculation of the parameters against ethanol were not statistically practical due to the low affinity.

| Alcohol substrates | $k_{\text{cat}}$ (1/s) | $K_M$ (mM) | $k_{\text{cat}}/K_M$ (1/M/s) |
| --- | --- | --- | --- |
| Ethanol* | $0.8 \pm 1.0$ | $1232.5 \pm 1986.3$ | $0.6 \pm 1.3$ |
| Butanol | $1.9 \pm 0.4$ | $195.7 \pm 87.3$ | $10.5 \pm 5.1$ |
| Isobutanol | $2.3 \pm 0.1$ | $180.1 \pm 21.7$ | $13.0 \pm 1.7$ |
| Prenol | $2.7 \pm 0.3$ | $101.0 \pm 21.6$ | $26.4 \pm 6.6$ |
| Furfuryl alcohol | $1.1 \pm 0.1$ | $37.1 \pm 9.8$ | $29.1 \pm 8.4$ |
| Pentanol | $3.9 \pm 1.0$ | $64.7 \pm 33.6$ | $59.1 \pm 19.3$ |
| Isoamyl alcohol | $2.8 \pm 0.3$ | $59.9 \pm 26.6$ | $46.0 \pm 11.9$ |
| Benzyl alcohol | $1.6 \pm 0.1$ | $12.2 \pm 1.6$ | $130.3 \pm 17.8$ |
| Hexanol | $6.4 \pm 0.4$ | $29.2 \pm 4.0$ | $219.5 \pm 33.5$ |
| 3-cis-Hexen-1-ol | $2.3 \pm 0.2$ | $6.4 \pm 2.2$ | $360.1 \pm 129.4$ |
| Phenylethyl alcohol | $6.1 \pm 0.1$ | $10.6 \pm 0.4$ | $577.1 \pm 26.2$ |
| 3-Methoxybenzyl alcohol | $18.8 \pm 0.9$ | $10.0 \pm 1.2$ | $1,891.6 \pm 248.8$ |
| Geraniol | $8.0 \pm 0.2$ | $4.1 \pm 0.4$ | $1,975.7 \pm 186.8$ |
| Citronellol | $7.7 \pm 0.2$ | $3.2 \pm 0.3$ | $2,400.9 \pm 247.9$ |
| Nerol | $2.2 \pm 0.04$ | $4.1 \pm 0.8$ | $4,119.5 \pm 753.0$ |
| Chloramphenicol | $105.5 \pm 6.9$ | $0.2 \pm 0.05$ | $491,901.5 \pm 115,286.2$ |

In addition, we found acetylation of fatty alcohols such as octanol and decanol to produce long-chain esters (Figs. S6B, S6C) that can potentially be used for drop-in biodiesel applications. The alcohol compatibility of CATec3 Y20F expanded from ethanol to terpenoid alcohols such as geraniol and nerol. Due to high K_M_ value (>1M) towards ethanol, CATec3 Y20F is more favorably applied for biosynthesis of higher-chain alcohol esters (Fig. S6A). This characteristic is potentially beneficial to produce designer esters rather than ethyl esters in organisms since ethanol is a common fermentative byproduct that can act as a competitive substrate. In comparison to CATsa Y20F A138T, CATec3 Y20F displayed higher activity towards not only isobutanol, but also most of other alcohols (Fig. S7). It is noteworthy that these engineered CATs exhibited different alcohol specificities (Table S3). For example, CATsa Y20F A138T was relatively more specific to phenylethyl alcohol than terpenoid alcohols as compared to CATec3 Y20F (Fig. S7).

We next examined whether CATec3 Y20F was also compatible with longer-chain acyl-CoAs. We analyzed the relative activities of CATec3 Y20F against a set of 10 linear, branched, and aromatic acyl-CoAs that can be synthesized in organisms together with isobutanol as a co-substrate (Figs. 2I). CATec3 Y20F has the highest activity towards the native substrate acetyl-CoA, which is the most abundant and critical precursor metabolite for cell biosynthesis. As compared to acetyl-CoA, CATec3 Y20F achieved 46%, 28%, 15%, 12%, 11%, and 9% of activity towards isobutyl-CoA, propionyl-CoA, butyryl-CoA, valeryl-CoA, phenylethyl-CoA, and isovaleryl-CoA, respectively. No activity was detected against linear fatty acyl-CoAs longer than valeryl-CoA (Fig. 2I). Interestingly, CATec3 Y20F also exhibited activity towards an uncommon lactyl-CoA for lactate ester biosynthesis. Since lactyl-CoA is not commercially available for *in vitro* assay, we determined the activity *in vivo* by using a recombinant *E. coli* co-expressing CATec3 Y20F and a propionyl-CoA transferase (PCT) derived from different microbes including *Thermus thermophilus* (PCTtt) that transfers CoA from acetyl-CoA to lactate (40, 41) (Fig. S8). By co-feeding the recombinant *E. coli* with 2 g/L of isoamyl alcohol and lactate, isoamyl lactate could be produced at a level of about 66.6 mg/L (Fig. S8B), which is at least 2.5-fold higher than the use of the eukaryotic AATs reported previously (9). Since PCTtt is derived from a thermophile, the lactate ester biosynthesis pathway is likely robust and compatible with thermophilic hosts. Taken together, we have shown unequivocally that CATs can be repurposed for *de novo* thermostable AATs (Fig. 2J). The engineered CATec3 Y20F exhibits extraordinary robustness, efficiency, and compatibility with various alcohols and acyl-CoA moieties, making it an ideal platform to synthesize designer bioesters in multiple organisms.

### Repurposing the engineered CAT for designer ester biosynthesis in a mesophilic *E. coli*

Microbial conversion presents a renewable and sustainable route to synthesize chemicals. Here, we characterized how efficient and compatible the engineered CATec3 Y20F is to enable biosynthesis of designer esters in a mesophilic *E. coli*, a workhorse for industrial biotechnology. To demonstrate the biosynthesis of designer acetate esters, we engineered HSEC01, a recombinant *E. coli* BL21 (DE3) harboring CATec3 Y20F under the control of a T7lac promoter that uses its native metabolism to convert fermentable sugars into the precursor acetyl-CoA, and characterized the engineered strain in the growth media supplemented with representative linear, branched, saturated, unsaturated, and/or aromatic alcohols and with hexadecane for *in situ* ester extraction (Fig. 3A). The result showed that the recombinant CATec3 Y20F-expressing *E. coli* could produce all of the expected acetate esters with relatively high efficiency and compatibility. In batch cultures, conversion of all alcohols to their respective acetate esters achieved more than 50% (mol/mol) yield within 24 h (Fig. 3B). Noticeably, yields of phenylethyl and geranyl acetate could reach up to 80% (mol/mol). The recombinant *E. coli* produced and secreted designer bioesters at final titers of 2.6 g/L butyl acetate, 2.3 g/L of isobutyl acetate, 3.1 g/L pentyl acetate, 2.9 g/L isoamyl acetate, 2.6 g/L 3-hexenyl acetate, 0.9 g/L benzyl acetate, 1.2 g/L 2-phenylethyl acetate, and 0.3 g/L geranyl acetate (Fig. S9). To further increase ester production, we performed fed-batch fermentation using the representative branched isoamyl alcohol and aromatic phenylethyl alcohol. With the intermittent feeding of up to 10 g/L alcohols, the recombinant *E. coli* could produce the expected esters at a relatively high efficiency, achieving titers of 13.9 g/L and 10.7 g/L and yields of 95% (mol/mol) and 80% (mol/mol) for isoamyl acetate and phenylethyl acetate, respectively (Figs. 3C). We did not observe any noticeable growth inhibition at this high level of ester production in the fed batch mode due to the *in situ* ester extraction with hexadecane (Fig. S10), even though both alcohols and esters are known to be toxic to microbial health at relatively low concentrations (< 2 g/L) (42, 43). As the heterologous pathways for higher alcohols have been metabolically engineered in recombinant organisms (e.g., *E. coli*) (34, 44–46), designer bioesters can be produced by using the engineered CAT and either co-feeding fermentable sugars and alcohols as demonstrated here or via natural fermentative processes which produce alcohols natively.

**Figure 3.**
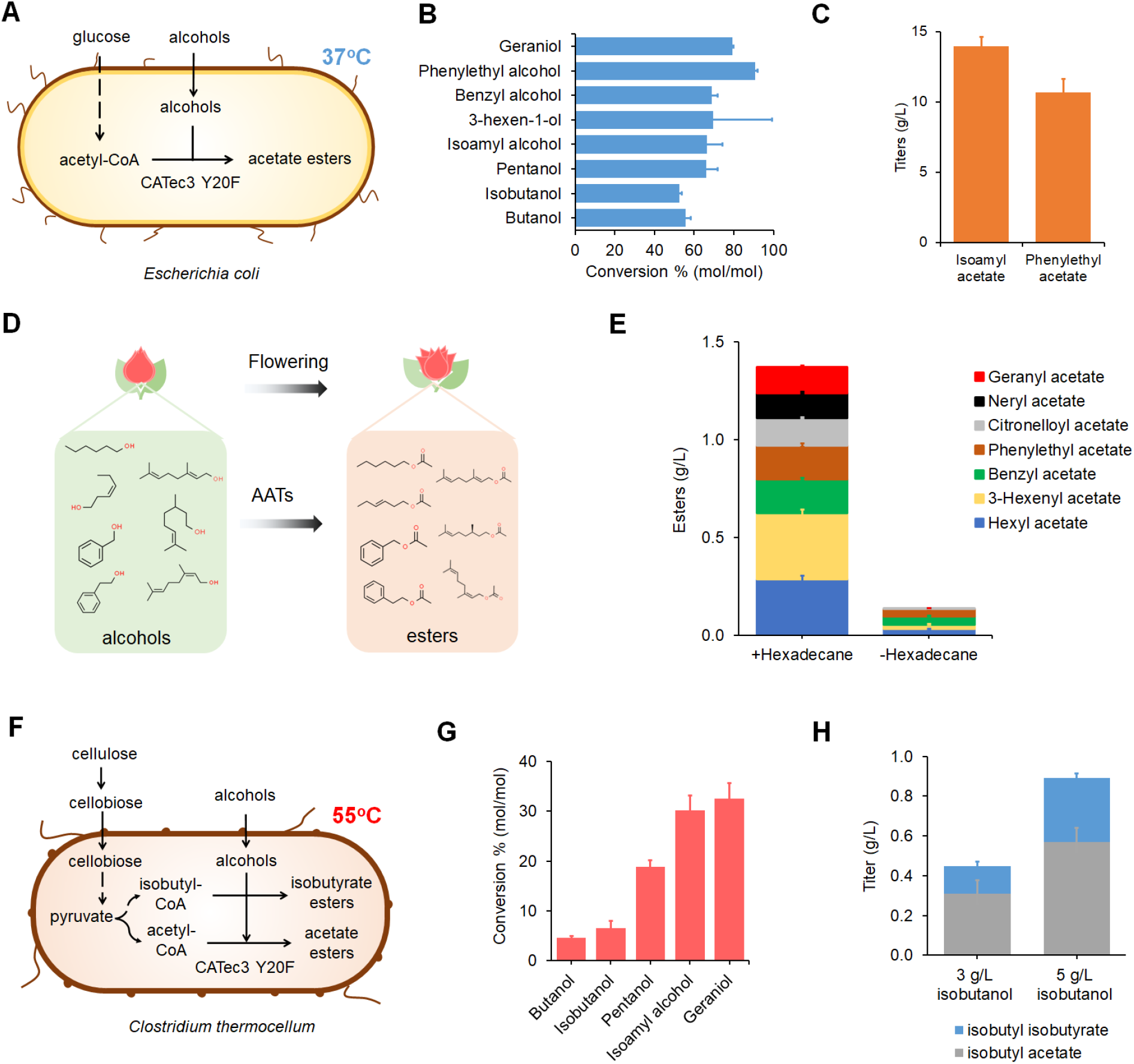
Ester biosynthesis by mesophilic and thermophilic organisms harboring the robust and efficient CATec3 Y20F. **(A)** Designer ester biosynthesis at 37°C by co-feeding a CATec3 Y20F-expressing *E. coli* strain with glucose and various alcohols. **(B)** Alcohol conversion efficiencies. Each alcohol was supplemented in the medium at 3 g/L for n-butanol, isobutanol, n-pentanol, and isoamyl alcohol, 1 g/L for benzyl alcohol and phenylethyl alcohol, and 0.3 g/L for geraniol. **(C)** Fed-batch conversion of isoamyl alcohol to isoamyl acetate by the CATec3 Y20F-expressing *E. coli* HSEC01. **(D)** Ester biosynthesis in developing rose petals. Roses *(Rosa hybrida)* synthesize volatile esters by alcohol acyltransferases (AATs), contributing to its unique aroma. **(E)** Simulated ester profile of roses by HSEC01. **(F)** Designer ester biosynthesis at 55°C by co-feeding a CATec3 Y20F-expressing *C. thermocellum* strain HSCT2108 with cellulose and various alcohols. **(G)** Alcohol conversion efficiencies. Each alcohol was supplemented in the medium at 3 g/L n-butanol, 3 g/L isobutanol, 3 g/L n-pentanol, 3 g/L isoamyl alcohol, and 0.3 g/L of geraniol. **(H)** Isobutyl ester production of HSCT2108 with different supplementation of isobutanol concentrations. In panels **B-C, E**, and **G-H**, each value represent mean ± 1 stdev from three biological replicates.

### Naturally reconstituting an ester profile of roses in *E. coli*

Due to high efficiency and compatibility of the engineered CAT for designer ester biosynthesis in the recombinant CATec3 Y20F-expressing *E. coli* HSEC01, we further explored whether CATec3 Y20F could be harnessed as an efficient AAT to create ester profiles exhibited by floral plants. In nature, eukaryotic AATs in floral plants formulate a mixture of volatile esters that contribute to their beneficial interactions with the environment. For example, roses *(Rosa hybrida)* utilizes AATs, regulated within their developing stages (47), to synthesize and emit volatile esters for attracting pollinators (Fig. 3D). To mimic an ester profile of *Rosa hybrida*, we fed HSEC01 with a mixture of alcohols at a total working concentration of 1 g/L, consisting of 0.2 g/L hexanol, 0.2 g/L 0.15 g/L 3-cis-hexen-1-ol, benzyl alcohol, 0.15 g/L phenylethyl alcohol, 0.1 g/L geraniol, 0.1 g/L nerol, and 0.1 g/L citronellol at a mid-log phase (OD_600nm_~1.0). The recombinant *E. coli* was capable of rapidly and completely converting the alcohol mixture into the desirable acetate ester profiles with a yield of 97.1 ± 0.7 % (mol/mol) and a titer of ~1.5 g/L within 12 h (Figs. 3E, S11A). For high production of esters, the *in situ* extraction using hexadecane overlay is very critical to mitigate the toxicity of esters (Fig. 3E, S11B).

### Enabling designer ester biosynthesis in a cellulolytic thermophile *C. thermocellum*

Given the native metabolic capability for effectively degrading recalcitrant cellulosic biomass at elevated temperatures (≥ 50°C), *C. thermocellum*, a cellulolytic, thermophilic, obligate anaerobic, gram-positive bacterium, has emerged as a microbial platform for consolidated bioprocessing (CBP) where cellulolytic enzyme synthesis, biomass hydrolysis, and fermentation can take place in a single step to produce desirable chemicals. By harnessing the high thermostability of CATec3 Y20F, we investigated whether CATec3 Y20F-harboring *C. thermocellum* could catabolize cellulose and convert various alcohols to designer bioesters efficiently at an elevated temperature of 55°C (Fig. 3F). We chose the engineered *C. thermocellum* Δ*clo1313_0613*, Δ*clo1313_0693* as the host because its two carbohydrate esterases was disrupted to alleviate ester degradation (30). By co-feeding cellulose and each higher alcohol, the recombinant *C. thermocellum* could produce all respective acetate esters (Fig. 3G, Fig. S12A). Since *C. thermocellum* has the endogenous isobutyl-CoA pathway(48), we also observed the production of isobutyrate esters such as butyl isobutyrate and isobutyl isobutyrate as byproducts (Fig. S12B). Many of these esters, such as n-butyl, n-pentyl, isoamyl, and geranyl esters, have never been reported to be feasibly synthesized in a thermophile. Among the esters, isoamyl acetate was produced at the highest conversion yield of > 30% (mol/mol) and titer of 1.2 g/L. Ester production in *C. thermocellum* was not as high as observed in *E. coli* likely due to the metabolic burden required to make cellulolytic enzymes for cellulose degradation along with overexpression of the heterologous gene. An increased titer of isobutyl esters was achieved when feeding a higher concentration of isobutanol (Figure 3H) below the lethal concentration (Figure S13), indicating that the enzyme expression and/or alcohol availability were likely unsaturated in *C. thermocellum*. Further strain engineering and/or optimization of medium and operating conditions might help boost the production of designer bioesters by the CBP technology.

### Thermostability of the engineered CAT played a critical role for efficient designer ester biosynthesis at elevated temperatures in *C. thermocellum*

Functional expression of a heterologous protein in thermophiles requires high thermostability, which is not well understood and often presents a significant bottleneck in metabolic engineering of these organisms. Inspired by the differences in the catalytic efficiency and melting temperatures among the engineered CATs (Table S3), we investigated how thermostability of the CATs affected ester production in *C. thermocellum*. We characterized the recombinant *C. thermocellum* Δ*clo1313_0613* Δ*clo1313_0693* harboring various CATs with distinctive melting temperatures and catalytic efficiency for *in vivo* isobutyl ester production by co-feeding cellulose and isobutanol at 55°C (Fig. 4A). Among the recombinant *C. thermocellum* strains, HSCT2108 harboring CATec3 Y20F, which has the highest catalytic efficiency and melting temperature, produced the highest level of isobutyl esters (892 mg/L), about 14-fold higher than the CATsa F97W-expressing *C. thermocellum* HSCT2105. Even though CATsa Y20F A138T has similar catalytic efficiency but higher melting temperature relative to CATsa Y20F, the CATsa Y20F A138T-expressing strain HSCT2113 produced 46% more esters than the CATsa Y20F-expressing strain HSCT2106. Similarly, we also observed higher ester production in the CATec3-expressing strain HSCT2107 than the CATsa F97W-expressing strain HSCT2105, where CATec3 has similar catalytic efficiency but higher melting temperature. Remarkably, both HSCT2107 and HSCT2113 produced esters at very similar level, although CATsa Y20F A138T has higher catalytic efficiency but about 10°C lower melting temperature than CATec3. These results strongly suggested that CAT robustness with enhanced thermostability plays a critical role for efficient ester production in *C. thermocellum* at elevated temperatures.

**Figure 4.**
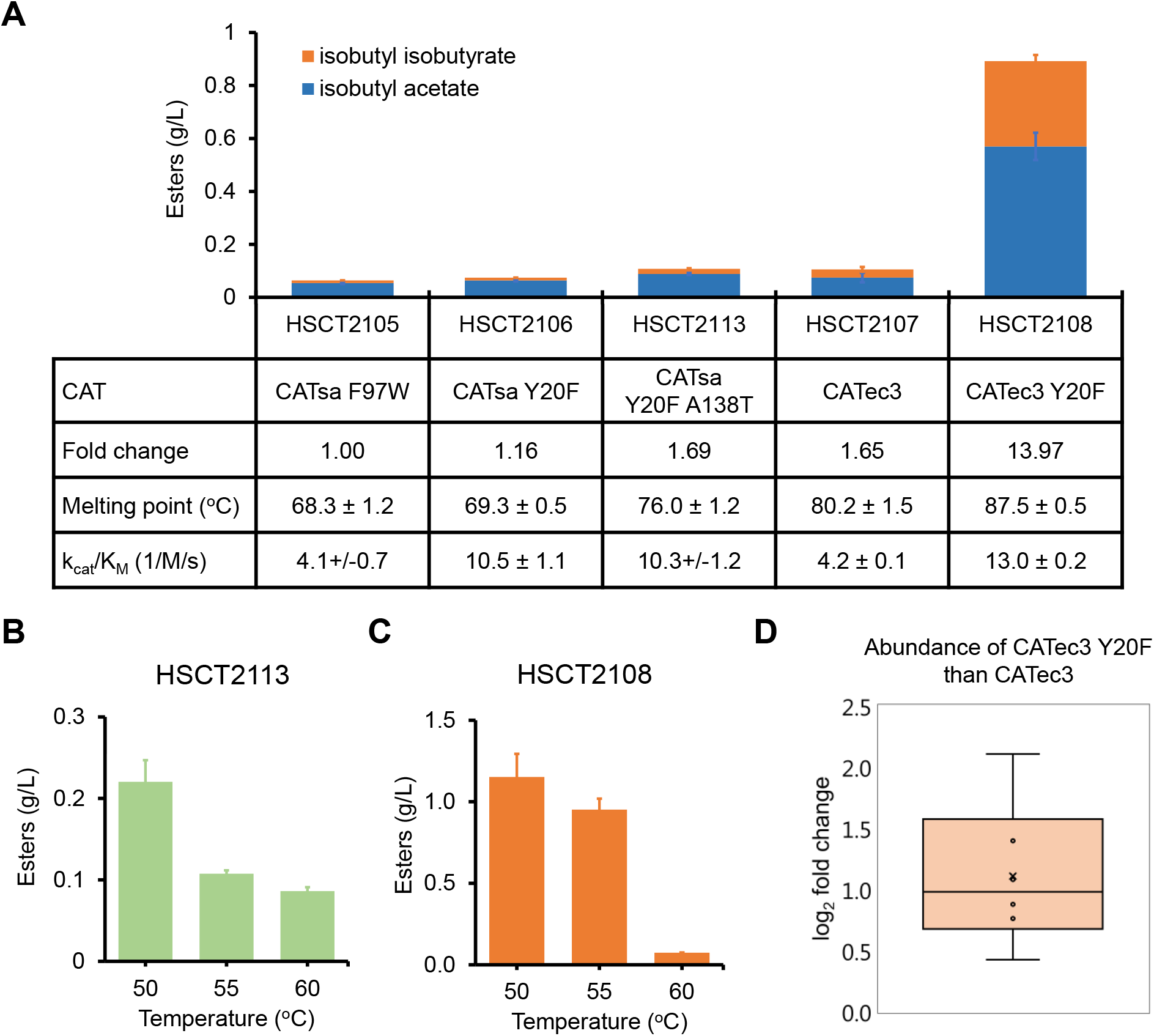
Effect of CAT thermostability on robust and efficient microbial biosynthesis of esters at elevated temperatures. **(A)** Isobutyl ester production of the recombinant *C. thermocellum* strains harboring various CATs. **(B-C)** Effect of temperatures on isobutyl ester production by the **(B)** CATec3-expressing strain HSCT2107 and **(C)** CATec3 Y20F-expressing strain HSCT2108. In panels **A-C**, each data represents mean ± 1 stdev from three biological replicates. Isobutanol (5 g/L) was fed at the early stationary phase of the culture to avoid cell growth inhibition (Fig. S13). **(D)** Comparison of CAT tryptic peptide abundance differences between HSCT2108 and HSCT2107 on a peptide-by-peptide basis (n = 6 peptides). The average (‘x’) and median (‘-’) log2 peptide differences were 1.1 and 1.0 respectively. The data is represented as quartiles from three biological replicates (p-value < 0.005, see also Fig. S14B).

To further elucidate the effect of thermostability of CATs on ester production, we characterized the performance of HSCT2113 and HSCT2108 at various elevated temperatures (Fig 4B, 4C) compatible with *C. thermocellum* growth. Interestingly, HSCT2113 increased the ester production up to 220 mg/L at 50°C, about 2-fold higher than 55°C (Fig. 4B). In contrast, HSCT2108 produced esters at relatively similar level of about 1 g/L at 50 and 55°C, while the production was reduced to 74 mg/L at 60°C (Fig. 4C). Proteolysis is a common cellular process to degrade and remove denatured or misfolded proteins (49). If the cell growth temperature affected integrity of the CATs, their intracellular abundances would be altered. To investigate changes in the intracellular abundance of CATs, we analyzed and compared proteomes across the two representative strains expressing CATec3, the wildtype (HSCT2107) versus the Y20F mutant (HSCT2108). Since the only difference between these two strains is one amino acid substitution, we could reliably quantify the relative abundance of each CAT by comparing tryptic peptide fragments (Fig. S14A). The result showed that CATec3 Y20F was 2.2-fold (average difference between peptide abundances) more abundant than CATec3 (Fig. 4D). Because CATec3 Y20F has a 7°C higher melting temperature than CATec3 (Table S3), the intracellular protein abundance change might imply a lesser degree of denaturation or misfolding due to higher protein thermostability. Taken altogether, CAT thermostability is critical for robust and efficient ester production in thermophiles by maintaining its intracellular protein abundance.

## Materials and Methods

### Protein homology modeling of CATs, *in silico* mutagenesis, and protein contact analysis

We used the Swiss-Model (50) and the ‘Builder’ tools of the commercial software MOE (Molecular Operating Environment software, version 2019.01) (51) to generate the three-dimensional (3D) structures of CATs, alcohols, acyl-CoAs, and their complexes, as previously described (29). To perform docking simulations in MOE, we started by searching the potential binding pocket with the ‘ Site Finder’ tool and selecting the best-scored site that is consistent with the reported catalytic sites (52). Next, we performed docking simulations for acyl-CoA and alcohol with CATs using the induced fit protocol with the Triangle Matcher placement method and the London ΔG scoring function. We selected the best-scored binding pose exhibiting the interaction between the residue and the substrate at root-mean-square-deviation (RMSD) < 2.3 Å. We used the ‘alanine scan’ and ‘residue scan’ tools of MOE to identify the potential residue candidates for mutagenesis of the acyl-CoA-alcohol-CAT complex, based on the ΔStability and/or ΔAffinity values calculated. Mutant candidates with small values of the ΔStability and/or ΔAffinity are chosen for experimental testing. To perform the protein contact analysis, we used the ‘Protein Contacts’ tool of MOE.

### Phylogenetic tree of CATs

Multiple sequence alignment was performed using MEGA7 (53) as previously described (29). Phylogenetic tree of CATs was built based on the aligned sequences using the maximum likelihood method with 1,000 bootstrap replicates. A 40% bootstrap confidence level cutoff was selected.

### Molecular cloning and site-directed mutagenesis

The primers used in this study are listed in Table S5. Plasmids were constructed by Gibson DNA assembly using the PCR products amplified by the primers. All the constructed plasmids were checked by PCR amplification and Sanger sequencing. Site-directed mutagenesis was performed using the QuikChange^™^ site-directed mutagenesis protocol using the Phusion DNA polymerase (Thermo Fisher Scientific, MA, USA) and the listed primers (54, 55).

### Enzyme characterization

To determine *in vitro* melting temperatures and catalytic efficiencies, His-tagged CATs were purified and characterized as previously described (29). In the 5,5’-dithiobis-2-nitrobenzoic acid (DTNB) assay, final enzyme concentrations of 0.05-0.1 μg/mL and 5-10 μg/mL were used for the reactions with chloramphenicol and alcohols, respectively. For heat inactivation experiments, 50 μL of the purified CATs were incubated at the temperatures from 50 to 70°C in a thermocycler for an hour, with the lid temperature set at 70°C. Residual activity was measured at 37°C using chloramphenicol and acetyl-CoA as substrates and normalized by the activity of the samples incubated at 50°C. To determine Michaelis-Menten kinetics, concentrations of the alcohol substrates varied as follows: (i) 0-400 mM for ethanol, butanol, and isobutanol, (ii) 0-100 mM for pentanol, isoamyl alcohol, 3-cis-hexen-1-ol, prenol, and furfuryl alcohol, (iii) 0-2 mM for octanol, (iv) 0-0.2 mM for decanol, (v) 0-50 mM for hexanol, citronellol, farnesol, and nerol, (vi) 0-40 mM for 3-methoxybenzyl alcohol, benzyl alcohol, and geraniol, and (vii) 0-20 mM 2-phenylethyl alcohol. For the alcohols with low solubility, 10% (w/v) DMSO was supplemented in the reaction solution. The enzyme reactions were held at 50°C in a BioTek microplate reader for at least 30 minutes with measurements every one minute. The kinetic parameters were calculated using the non-linear regression method as previously described (29).

### Strains and plasmids

*E. coli* BL21(DE3) was used for protein expression and purification, and alcohol conversion experiments. Plasmids used in this study were listed in Table S6. For *C. thermocellum* experiments, the strain HSCT2005 lacking Clo1313_0613 and Clo1313_0693 was used (30).

### Media and cell cultivation

*E. coli* strains were grown in lysogeny broth (LB) medium or M9 hybrid medium containing glucose as a carbon source and 5 g/L yeast extract supplemented with 100 μg/mL ampicillin and/or 50 μg/mL kanamycin when appropriate. *C. thermocellum* strains were cultured in an anaerobic chamber (Sheldon manufacturing, OR, USA) with an anaerobic gas mixture (90% N_2_, 5% CO_2_, 5% H_2_) or rubber stopper sealed anaerobic Balch tubes outside the chamber. For *C. thermocellum* transformation, CTFuD or CTFuD-NY (56) media was used. The CTFuD medium contained 2 g/L yeast extract while CTFuD-NY used vitamins and trace elements instead of the yeast extract. To maintain the plasmids in *C. thermocellum*, 10 μg/mL thiamphenicol was supplemented. For alcohol conversion experiments with *C. thermocellum*, strains were grown in a defined C-MTC medium as previously described (30, 57)

### *C. thermocellum* transformation

*C. thermocellum* cells were transformed by electroporation as previously described (29, 56). A series of two consecutive exponential pulses were applied using the electroporation system (cat # 45-0651, BTX Technologies Inc., MA, USA) set at 1.8 kV, 25 μF, and 350 Ω, which usually resulted in a pulse duration of 7.0-8.0 ms.

### *In vivo* screening of CATSa variants

To prepare pre-cultures, single colonies from LB agar plates were first inoculated into 100 μL of LB in 96-well microplates using sterile pipette tips. The precultures were then grown overnight at 37°C and 400 rpm in an incubating microplate shaker (Fisher Scientific, PA, USA). Next, 5 % (v/v) of pre-cultures were inoculated into 100 μL of the M9 hybrid media containing 20 g/L of glucose, 0.1 mM of IPTG, and 2 g/L of isobutanol in a 96-well microplate with hexadecane overlay, containing isoamyl alcohol as an internal standard, in a 1:1 (v/v) ratio. The microplates were sealed with a plastic adhesive sealing film, SealPlate^®^ (EXCEL Scientific, Inc., CA, USA) and incubated at 37°C and 400 rpm for 24 h in an incubating microplate shaker. Samples from the hexadecane layer were collected and subjected to GC/MS for ester identification and quantification.

### Conversion of alcohols to esters in *E. coli*

For batch cultures, tube-scale alcohol conversions were performed in 4 mL M9 medium containing 10 g/L glucose with addition of 1 mL of hexadecane for *in situ* extraction at 37°C. 0.1 mM isopropyl β-D-1-thiogalactopyranoside (IPTG) was initially added to induce expression of CATec3 Y20F. Alcohols were supplemented in the initial medium, and the product yield and titer were measured at 12h, 24h, and 48h time points. For fedbatch cultures designed to achieve high-level conversion of alcohols (i.e., isoamyl alcohol, phenylethyl alcohol), cells were grown microaerobically in a 125 mL screw capped shake flask with a working volume of 20 mL M9 medium containing 25 g/L glucose and 10 mL hexadecane. A volume of 25-50 μL of the alcohols (? 98% purity) were added to the culture at 6h, 9h, 12h, 15h, and 24h time points with a working concentration of 2 g/L per addition.

### Conversion of alcohols to esters in *C. thermocellum*

Tube-scale cellulose fermentation was performed in the batch mode as previously described (29). Briefly, 19 g/L of Avicel PH-101 was used as a sole carbon source in a 16 mL culture volume. 0.8 mL of overnight cell culture was inoculated in 15.2 mL of C-MTC medium, and 4 mL hexadecane was added in the anaerobic chamber. Each tube contained a small magnetic stirrer bar to homogenize cellulose, and the culture was incubated in a water bath connected with a temperature controller and a magnetic stirring system. Alcohols were fed to the culture at 36h time point when cells entered early stationary growth phase. pH was adjusted to between 6.4 and 7.8 with 5 M KOH injection.

### High-performance liquid chromatography (HPLC) analysis

HPLC system (Shimadzu Inc., MD, USA) was used to quantify extracellular metabolites. 800 μL of samples were centrifuged at 17,000 x g for 3 minutes followed by filtering through 0.2 micron filters. The samples were run with 5 mM H_2_SO_4_ at 0.6 mL/min on an Aminex HPX-87H (Biorad Inc., CA, USA) column at 50°C. Refractive index detector (RID) and ultra-violet detector (UVD) at 220 nm were used to determine concentrations of sugars, organic acids, and alcohols.

### Gas chromatography coupled with mass spectroscopy (GC/MS) analysis

GC (HP 6890, Agilent, CA, USA) equipped with a MS (HP 5973, Agilent, CA, USA) was used to quantify esters. A Zebron ZB-5 (Phenomenex, CA, USA) capillary column (30 m x 0.25 mm x 0.25 μm) was used with helium as the carrier gas at a flow rate of 0.5 mL/min. The oven temperature program was set as follows: 50°C initial temperature, 1°C/min ramp up to 58°C, 25°C/min ramp up to 235°C, 50°C/min ramp up to 300°C, and 2-minutes bake-out at 300°C. 1 μL sample was injected into the column with the splitless mode at an injector temperature of 280°C. For the MS system, selected ion mode (SIM) was used to detect and quantify esters. As an internal standard, 10 mg/L n-decane were added in initial hexadecane layer and detected with m/z 85, 99, and 113 from 12 to 15 minute retention time range.

### Proteomics

For proteomics, cell cultures were sampled at 48h in the middle of stationary growth phase and stored in −80°C before the analysis. Two milliliters of a slurry containing *C. thermocellum* cells growing on Avicel was pelleted by centrifugation (5000 x g for 10 min), supernatant discarded, and pellet resuspended in 350 μl of lysis buffer (4% SDS, 5 mM DTT, 100 mM Tris-HCl, pH 8.0). Cells were lysed by sonication (Branson Sonifier; 20% amplitude, 2 s pulses, 30 s total), precleared by centrifugation (21000 x g for 10 min) and crude protein supernatant with a Nanodrop OneC spectrophotometer (Thermo Scientific). Samples were then adjusted 15 mM iodoacetamide (20 min at room temperature in the dark) and 300 μg cleaned up via protein aggregation capture (58). Aggregated protein (on 1 micron magnetic Sera-Mag beads; GE Healthcare) was then digested with proteomics-grade trypsin (1:75 w/w; Promega) in 100 mM Tris-HCl, pH 8.0 overnight at 37°C, and again for 3 h at 37°C the following day. Released tryptic peptides were then acidified to 0.5% formic acid, filtered through a 10 kDa MWCO spin filter (Vivaspin 2; Sartorius), and quantified by Nanodrop OneC. Three micrograms of peptides were then analyzed by 1D LC-MS/MS using a Vanquish uHPLC coupled directly to an Orbitrap Q Exactive mass spectrometer (Thermo Scientific) as previously described (59). Peptides were separated across a 180 min organic gradient using an in-house pulled nanospray emitter packed with 15 cm of 1.7-micron Kinetex reversed-phase resin (Phenomenex). Peptide fragmentation spectra were analyzed/sequenced by Proteome Discoverer software (Thermo Scientific) and peptides quantified by chromatographic area-under-the-curve. Peptide abundances were log2 transformed, distributions normalized by LOESS, and rolled up to their respective proteins via RRollup InfernoRDN (60). Protein abundance distributions were then median centered and statistical analyses performed in Perseus (61). Manual determination of CATec3 abundance across strains was performed by comparing log2 peptide abundance trends across orthologs as only one AA difference differentiate the two protein forms. All LC-MS/MS raw data have been deposited into the MassIVE and ProteomeXchange repositories with the following accession numbers: MSV000086201 (MassIVE) and PXD021695 (ProteomeXchange). Data can be downloaded directly via ftp://massive.ucsd.edu/MSV000086201/

## Conclusions

Esters are industrially important chemicals with applications as flavors, fragrances, solvents, and drop-in fuels. Robust and efficient AATs are an essential module to expand biological routes for sustainable and renewable production of designer bioesters. Through bioprospecting and rational protein engineering, we repurposed CATs to function as robust and efficient AATs that exhibit high compatibility to a broad range of pathways and microbial hosts. We demonstrated that the repurposed CATs are capable of producing designer esters in both mesophilic and thermophilic microorganisms with high efficiency, robustness, and compatibility. Using proteomics and comparative analysis of the engineered CATs, we found that the CAT robustness with enhanced thermostability is critical for efficient ester production in thermophiles by maintaining high level of intracellular CAT abundance. This work not only presents a robust, efficient, and highly compatible AAT platform for designer bioester production, but also elucidates the impact of enzyme thermostability on engineering heterologous pathways in a thermophilic whole-cell biocatalyst.

## Acknowledgements

This research was financially supported by the NSF CAREER award (NSF#1553250) and the Center for Bioenergy Innovation (CBI), the U.S. Department of Energy (DOE) Bioenergy Research Centers Funded by the Office of Biological and Environmental Research in the DOE Office of Science. The authors would like to acknowledge the Center of Environmental Biotechnology at UTK for using the GC/MS instrument, and the Joint Genome Institute (JGI) for gene synthesis.

