## Supplementary Materials for "Repurposing chloramphenicol acetyltransferase for a robust and efficient designer ester biosynthesis platform"

### Supplemental Tables

**Table S1.** Screening of CATsa variants for isobutyl ester synthesis in *E. coli*. Ethyl ester is a byproduct since *E. coli* produced ethanol endogenously. Each value represents mean ± 1 standard deviation (stdev) from at least three biological replicates.

| CATsa variants | Ethyl acetate (mg/L) | Isobutyl acetate (mg/L) |
|---|---|---|
| WT | 2.88 ± 0.24 | 4.16 ± 1.23 |
| Y20W | 2.12 ± 0.30 | 5.25 ± 3.51 |
| Y20H | 1.98 ± 0.45 | 3.75 ± 1.56 |
| G194K | 1.83 ± 0.36 | 1.33 ± 1.12 |
| Y20D | 1.98 ± 0.44 | 1.71 ± 1.44 |
| T88Y | 1.72 ± 0.17 | 1.72 ± 1.03 |
| T88D | 2.52 ± 0.83 | 1.81 ± 1.54 |
| T88K | 2.16 ± 0.66 | 1.14 ± 0.61 |
| Y20T | 1.92 ± 0.29 | 4.07 ± 0.95 |
| **Y20F** | 1.56 ± 0.15 | 181.80 ± 32.38 |
| G194R | 2.07 ± 0.59 | 4.09 ± 3.89 |
| T88L | 1.95 ± 0.16 | 0.58 ± 0.23 |
| G194Y | 2.06 ± 0.24 | 2.30 ± 0.69 |
| G194V | 2.03 ± 1.29 | 2.85 ± 2.47 |
| Y20L | 2.00 ± 0.45 | 12.88 ± 1.17 |
| T88M | 2.33 ± 0.83 | 3.52 ± 2.00 |
| L116R | 2.30 ± 0.75 | 7.99 ± 1.39 |
| L116K | 2.36 ± 0.11 | 12.62 ± 4.38 |
| F49Y | 2.13 ± 0.39 | 4.51 ± 2.92 |
| F49W | 2.28 ± 0.20 | 3.04 ± 2.60 |
| G92R | 2.75 ± 0.87 | 3.66 ± 2.50 |
| K48R | 2.20 ± 0.35 | 6.01 ± 0.86 |
| Y112K | 2.40 ± 0.52 | 2.85 ± 2.03 |
| K48I | 2.65 ± 1.36 | 9.11 ± 3.64 |
| T88I | 2.86 ± 0.72 | 3.43 ± 1.04 |
| T88R | 2.92 ± 1.23 | 4.54 ± 1.71 |
| S140L | 2.60 ± 1.08 | 4.46 ± 0.84 |

**Table S2.** Melting temperatures of a set of 28 chloramphenicol acetyltransferases (CATs). Each value represents mean ± 1 stdev. Note that CAT1_ECOLX and CAT3_ECOLX are also denoted as CATec1 and CATec3, respectively.

| No | Type | CATs | Origins | Uniprot (or NCBI) protein entry | $T_m$ (°C) | Replicates |
|---|---|---|---|---|---|---|
| 1 | A | CATsa | *Staphylococcus aureus* | WP_001010387.1 | 71.2 ± 1.1 | 6 |
| 2 | A | CAT_STRAC | *Streptomyces acrimycini* | P20074 | 77.5 ± 4.3 | 4 |
| 3 | A | CAT2_ECOLX | *Escherichia coli* | P22615 | 81.6 ± 0.8 | 11 |
| 4 | A | CAT_THEACI | *Thrmoactinomyces sp.* | WP_037996077.1 | 81.0 ± 0.0 | 2 |
| 5 | A | CAT_BACI | *Bacillus sp.* | A0A3N9QXS5 | 70.7 ± 0.9 | 3 |
| 6 | A | CAT4_STAAU | *Staphylococcus aureus* | P36882 | 71.0 ± 0.0 | 2 |
| 7 | A | CAT_BAOCI | *Bacillus oceanisediminis* | A0A160M771 | 65.0 ± 0.0 | 3 |
| 8 | A | CAT_CLOBU | *Clostridium butyricum* | Q02736 | 70.0 ± 0.0 | 3 |
| 9 | A | CAT3_STAAU | *Staphylococcus aureus* | P06135 | 68.7 ± 2.4 | 3 |
| 10 | A | CAT5_STAAU | *Staphylococcus aureus* | P36883 | 82.0 ± 0.0 | 2 |
| 11 | A | CAT_BACPU | *Bacillus pumilus* | P00487 | 72.0 ± 0.0 | 2 |
| 12 | A | CAT1_ECOLX | *Escherichia coli* | P62577 | 71.2 ± 0.7 | 6 |
| 13 | A | CAT_KLEPS | *Klebsiella sp.* | P58777 | 67.7 ± 0.9 | 3 |
| 14 | A | CAT_GEO | *Geomicrobium sp.* | WP_042419141.1 | 43.3 ± 0.8 | 4 |
| 15 | B | CAT4_ECOLX | *Escherichia coli* | P26838 | 71.8 ± 1.3 | 12 |
| 16 | B | CAT4_PSEAE | *Pseudomonas aeruginosa* | P26841 | 71.0 ± 1.6 | 4 |
| 17 | B | CAT4_MORMO | *Morganella morganii* | P50869 | 57.0 ± 0.0 | 2 |
| 18 | A | CAT_LYSI | *Lysinibacillus boronitolerans* | A0A0A3IEC4 | 76.0 ± 0.0 | 2 |
| 19 | A | CAT_VIBAN | *Vibrio anguillarum* | P49417 | 71.5 ± 0.5 | 2 |
| 20 | A | CAT1_CLOPF | *Clostridium perfringens* | P26825 | 65.0 ± 0.0 | 2 |
| 21 | A | CAT_PROMI | *Proteus mirabilis* | P07641 | 81.0 ± 0.0 | 2 |
| 22 | A | CAT3_ECOLX | *Escherichia coli* | P00484 | 80.2 ± 1.5 | 6 |
| 23 | B | CAT4_KLEAE | *Klebsiella aerogenes* | P50868 | 70.5 ± 0.9 | 4 |
| 24 | A | CAT_STRAG | *Streptococcus agalactiae* | Q03058 | 66.3 ± 5.8 | 4 |
| 25 | A | CAT2_HAEIF | *Haemophilus influenzae* | P22616 | 82.0 ± 1.8 | 6 |
| 26 | A | CAT_CAMCO | *Campylobacter coli* | P22782 | 69.0 ± 0.0 | 3 |
| 27 | A | CAT_CLODI | *Clostridioides difficile* | P11504 | n.a. | 2 |
| 28 | A | CAT_STAIN | *Staphylococcus intermedius* | P25309 | n.a. | 2 |

**Table S3.** Melting temperatures and kinetic parameters of CAT variants towards isobutanol. The co-substrate, acetyl-CoA, was supplemented at the saturated concentration of 2 mM. Kinetic parameters were determined at 50°C. Each value represents mean ± 1 stdev from at least three biological replicates.

| **CATs** | **Tm (°C)** | **$k_{cat}$ (1/s)** | **$K_M$ (mM)** | **$k_{cat}/K_M$ (1/M/s)** |
|---|---|---|---|---|
| CATec1 | 71.2 ± 0.7 | 0.4 ± 0.1 | 375.9 ± 121.7 | 1.0 ± 0.4 |
| CATec1 F102W | 68.0 ± 0.0 | 1.2 ± 0.1 | 259.3 ± 59.2 | 4.7 ± 1.2 |
| CATsa | 71.2 ± 1.1 | 0.3 ± 0.05 | 164.1 ± 29.6 | 2.1 ± 0.4 |
| CATsa F97W | 68.3 ± 1.2 | 0.6 ± 0.05 | 144.8 ± 23.7 | 4.1 ± 0.7 |
| CATsa Y20F | 69.3 ± 0.5 | 1.5 ± 0.06 | 146.3 ± 13.8 | 10.5 ± 1.1 |
| CATsa Y20F F97W | 68.7 ± 0.5 | 1.4 ± 0.1 | 212.3 ± 32.4 | 6.4 ± 1.1 |
| CATsa Y20F A138T | 76.0 ± 1.2 | 2.0 ± 0.09 | 189.2 ± 19.8 | 10.3 ± 1.2 |
| CATsa Y20F F97W A138T | 73.0 ± 0.6 | 1.4 ± 0.06 | 205.7 ± 17.8 | 6.7 ± 0.6 |
| CATec3 | 80.2 ± 1.5 | 0.7 ± 0.1 | 171.4 ± 24.6 | 4.2 ± 0.1 |
| CATec3 F97W | 81.2 ± 0.4 | 0.5 ±0.1 | 250.2 ± 84.3 | 1.8 ± 0.1 |
| CATec3 Y20F | 87.5 ± 0.5 | 2.3 ± 0.1 | 180.1 ± 21.7 | 13.0 ± 0.2 |
| CATec3 Y20F F97W | 85.5 ± 0.5 | 0.53 ± 0.1 | 156.7 ± 58.5 | 3.4 ± 0.1 |

**Table S4.** Kinetic parameters of CATsa Y20F A138T towards various alcohol substrates. The co-substrate, acetyl-CoA, was supplemented at the saturated concentration of 2 mM. The reactions were performed at 50°C. Each value represents mean ± 1 stdev from at least three biological replicates.

| **Alcohol substrates** | **$k_{cat}$ (1/s)** | **$K_M$ (mM)** | **$k_{cat}/K_M$ (1/M/s)** |
|---|---|---|---|
| Isobutanol | 2.0 ± 0.1 | 189.2 ± 19.8 | 10.3 ± 1.2 |
| Prenol | 2.9 ± 0.3 | 277.7 ± 33.9 | 10.3 ± 1.6 |
| Furfuryl alcohol | 0.9 ± 0.05 | 120.5 ± 10.2 | 7.6 ± 0.8 |
| Isoamyl alcohol | 10.7 ± 2.7 | 641.8 ± 204.3 | 16.6 ± 6.8 |
| 3-cis-hexen-1-ol | 1.7 ± 0.09 | 54.2 ± 4.5 | 32.1 ± 3.1 |
| 3-methoxybenzyl alcohol | 2.0 ± 0.1 | 39.2 ± 4.7 | 51.6 ± 7.2 |
| Geraniol | 1.9 ± 0.07 | 5.8 ± 0.7 | 325.5 ± 40.5 |
| Citronellol | 1.2 ± 0.04 | 2.7 ± 0.4 | 438.5 ± 64.6 |
| Nerol | 2.1 ± 0.06 | 2.6 ± 0.3 | 805.3 ± 106.3 |
| Chloramphenicol | 17.2 ± 0.5 | 0.3 ± 0.05 | 52,624.6 ± 7,524.1 |

**Table S5.** A list of primers used in this study. The underlines indicate mutation sites.

| Primer name | Primer sequence (5' to 3') |
|---|---|
| ***CATsa mutagenesis*** | |
| Y20W F | TTAATCAT TGG TTGAACCAACAAACGAC |
| Y20W R | GTTCAA CCA ATGATTAAATATCTCTTTTCTC |
| Y20H F | TTAATCAT CAT TTGAACCAACAAACGAC |
| Y20H R | GTTCAA ATG ATGATTAAATATCTCTTTTCTC |
| G194K F | GTTTGTGAT AAA TATCATGCAGGATTG |
| G194K R | TGCATGATA TTT ATCACAAACAGAATGATG |
| Y20D F | TTAATCAT GAT TTGAACCAACAAACGAC |
| Y20D R | GTTCAA ATC ATGATTAAATATCTCTTTTCTC |
| T88Y F | ACTTTAT TAT ATTTTTGATGGTGTATC |
| T88Y R | CAAAAAT ATA ATAAAGTGGCTCTAAC |
| T88D F | ACTTTAT GAT ATTTTTGATGGTGTATC |
| T88D R | CAAAAAT ATC ATAAAGTGGCTCTAAC |
| T88F F | ACTTTAT TTT ATTTTTGATGGTGTATC |
| T88F R | CAAAAAT AAA ATAAAGTGGCTCTAAC |
| Y20T F | TTAATCAT ACC TTGAACCAACAAACGAC |
| Y20T R | GTTCAA GGT ATGATTAAATATCTCTTTTCTC |
| Y20F F | TTAATCAT TTT TTGAACCAACAAACGAC |
| Y20F R | GTTCAA AAA ATGATTAAATATCTCTTTTCTC |
| G194R F | TGTGAT CGT TATCATGCAGGATTG |
| G194R R | TGATAA ACG TCACAAACAGAATGATGTA |
| T88L F | ACTTTAT CTG ATTTTTGATGGTGTATC |
| T88L R | CAAAAAT CAG ATAAAGTGGCTCTAAC |
| G194Y F | TGTGAT TAT TATCATGCAGGATTG |
| G194Y R | TGATAA ATA TCACAAACAGAATGATGTA |
| G194V F | TGTGAT GTT TATCATGCAGGATTG |
| G194V R | TGATAA AAC TCACAAACAGAATGATGTA |
| Y20L F | TTAATCAT CTG TTGAACCAACAAACGAC |
| Y20L R | GTTCAA CAG ATGATTAAATATCTCTTTTCTC |
| T88M F | ACTTTAT ATG ATTTTTGATGGTGTATC |
| T88M R | CAAAAAT CAT ATAAAGTGGCTCTAAC |
| L116R F | TATAC CGT TCTGATGTAGAGAAATA |
| L116R R | ATCAGA ACG GTATAAATCATAAAAC |
| L116K F | TATAC AAA TCTGATGTAGAGAAATA |
| L116K R | ATCAGA TTT GTATAAATCATAAAAC |
| F49Y F | CATTTAT TAT CTTAGTGACAAGGGTG |
| F49Y R | CTAAG ATA ATAAATGCAGGGTAAAATT |
| F49W F | CATTTAT TGG CTTAGTGACAAGGGTG |
| F49W R | CTAAG CCA ATAAATGCAGGGTAAAATT |
| G92R F | TTTGAT CGT GTATCTAAAACATTCTCTGG |
| G92R R | GATACA ACG TCAAAAATTGTATAAAGTGG |
| K48R F | GCATT CGT TTTCTTAGTGACAAGGGTGA |
| K48R R | AGAAA ACG AATGCAGGGTAAAATTTATATCC |
| Y112K F | AGTTT AAA GATTTATACCTTTCTG |

| | |
|---|---|
| Y112K R | AATC TTT AAACTCTTTGAAGTCAT |
| K48I F | GCATT AAT TTTCTTAGTGACAAGGGTGA |
| K48I R | AGAAA ATT AATGCAGGGTAAAATTTATATCC |
| T88I F | ACTTTAT ATT ATTTTTGATGGTGTATC |
| T88I R | CAAAAAT AAT ATAAAGTGGCTCTAAC |
| T88R F | ACTTTAT CGT ATTTTTGATGGTGTATC |
| T88R R | CAAAAAT ACG ATAAAGTGGCTCTAAC |
| S140L F | CTTTT CTG CTTTCTATTATTCCATGGAC |
| S140L R | AGAAAG CAG AAAAGCATTTTCAGGTATAGG |
| A138T F | GAAAAT ACC TTTTCTCTTTCTATTATTC |
| A138T R | AGAAAA GGT ATTTTCAGGTATAGGTG |
| ***CATec3 mutagenesis*** | |
| ec3 F97W F | AGACG TGG AGCGCGTTATCGTGCC |
| ec3 F97W R | GCGCT CCA CGTCTCTGTTTCCTGATG |
| ec3 Y20F F | AATTT TTT CGCCACAGACTGCCAT |
| ec3 Y20F R | TGGCG AAA AAATTCGAAATGTTCG |
| ***CATec1 mutagenesis*** | |
| ec1 F102W F | AAACC TGG AGCTCATTATGGAGTGAA |
| ec1 F102W R | GAGCT CCA GGTTTCCGTTTGTTC |
| ***CATec3 cloning under the control of PgapDH promoter*** | |
| PgapDH_ec3 F | TACAATAGGAGGCGATATTA ATGAATTATACTAAATTCGATGTC |
| PgapDH_ec3 R | AGAAACCTTTGTATATTTT TTACTTCAATTTCGAATTGCAGAG |
| **pHS0024 backbone PCR** | |
| HS216 | AAAATATACAAAGGTTTCTTGTG |
| HS498 | TAATATCGCCTCCTATTGTAAATTAAAAT |

**Table S6.** List of plasmids and strains used in this study.

| Name | Descriptions | Source |
|---|---|---|
| ***Plasmids*** | | |
| pET_CATsa | CATsa wild type encoding gene under the control of T7lac promoter, 6X His-tag at N-terminus | (1) |
| pET_CATsa Y20F A138T | CATsa Y20F A138T variant generated by site-directed mutagenesis from pET_CATsa | This study |
| pET_CATec3 | CAT3_ECOLIX (CATec3) encoding gene under the control of T7 promoter, 6X His-tag at C-terminus | This study |
| pET_CATec3 Y20F | CATec3 Y20F variant generated by site-directed mutagenesis from pET_CATec3 | This study |
| pHS0024 | pNW33N derivative, CATsa wild type gene under *C. thermocellum* gapDH promoter | (1) |
| pHS0024_F97W | pNW33N derivative, CATsa F97W under *C. thermocellum* gapDH promoter | (1) |
| pHS0024_Y20F | pNW33N derivative, CATsa Y20F under *C. thermocellum* gapDH promoter | This study |
| pHS0024_Y20F A138T | pNW33N derivative, CATsa_Y20F A138T under *C. thermocellum* gapDH promoter | This study |
| pHS0059 | pHS0024 harboring CATec3 replacing with CATsa | This study |
| pHS0059_Y20F | CATec3 Y20F under *C. thermocellum* gapDH promoter | This study |
| ***E. coli*** | | |
| BL21(DE3) | F– ompT gal dcm lon hsdSB($rB^-mB^-$) λ(DE3 [lacI lacUV5-T7p07 ind1 sam7 nin5]) [$malB^+$]K-12(λS) | Novagen |
| HSEC01 | BL21(DE3) harboring pET_CATec3 Y20F | This study |
| ***C. thermocellum*** | | |
| M1354 | *C. thermocellum* DSM1313 Δ*hpt* | (2) |
| HSCT2005 | *C. thermocellum* DSM1313 Δ*hpt* Δ*Clo1313_0613* Δ*Clo1313_0693* | (3) |
| HSCT2105 | HSCT2005 harboring pHS0024_F97W | (3) |
| HSCT2106 | HSCT2005 harboring pHS0024_Y20F | This study |
| HSCT2107 | HSCT2005 harboring pHS0059 | This study |
| HSCT2108 | HSCT2005 harboring pHS0059_Y20F | This study |
| HSCT2113 | HSCT2005 harboring pHS0024_Y20F A138T | This study |

### Supplemental Figures

**Figure S1.** *In silico* mutagenesis CATsa for enhanced affinity of isobutanol-acetyl-CoA-CATsa complex. **(A)** Docking simulation of isobutanol and acetyl-CoA on CATsa binding pocket. Residues close to the posed isobutanol and acetyl-CoA are shown. **(B)** Structural location of Tyr-20 and Phe-97 along with the catalytic active site, His-189 of CATsa. **(C-D)** Residue scan of residues interacting with isobutanol of the $CAT_{Sa}$-isobutanol-acetyl-CoA complex. Selected candidates are based on the **(C)** ΔAffinity values and **(D)** ΔAffinity and ΔStability (cutoff = 0 kcal/mol). **(E-F)** Residue scan of residues interacting with acetyl-CoA of the $CAT_{Sa}$-isobutanol-acetyl-CoA complex. Selected candidates are based on the **(E)** ΔAffinity values and **(F)** ΔAffinity and ΔStability (cutoff = -0.2 kcal/mol).

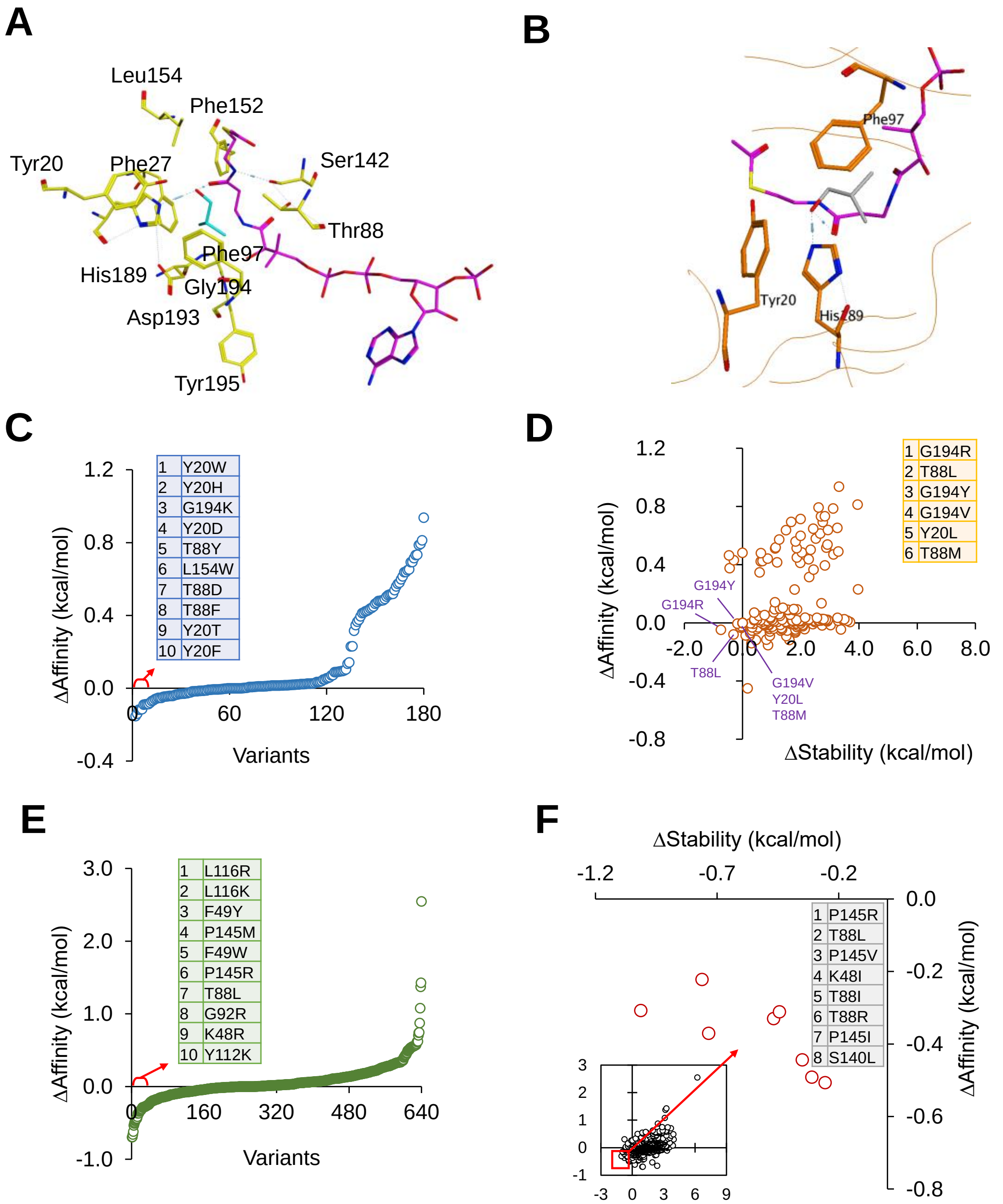

**Figure S2.** Screening of CAT activities towards isobutanol. **(A)** SDS-PAGE of His-tag purified CATs. The protein samples were loaded at 500 ng per lane. From 1 to 27; CATsa, CAT_STRAC, CAT2_ECOLX, CAT_THEACI, CAT_BACI, CAT4_STAAU, CAT_BAOCI, CAT3_STAAU, CAT_PROMI, CAT_BACPU, CAT1_ECOLIX, CAT_KLEPS, CAT_CLOBU, CAT_GEO, CAT_CAMCO, CAT4_ECOLIX, CAT4_PSEAE, CAT_CLODI, CAT_LYSI, CAT4_MORMO, CAT1_CLOPF, CAT5_STAAU, CAT3_ECOLX, CAT4_KLEAE, CAT_STRAG, CAT2_HAEIF, and CAT_VIBAN. The red arrow indicates the average size of the CATs (24.9kDa). CAT_STAIN was not included because of unsuccessful expression. **(B)** Specific activity of CATs towards 100 mM isobutanol at 50ºC. Each value represents mean ± 1 stdev from two biological replicates.

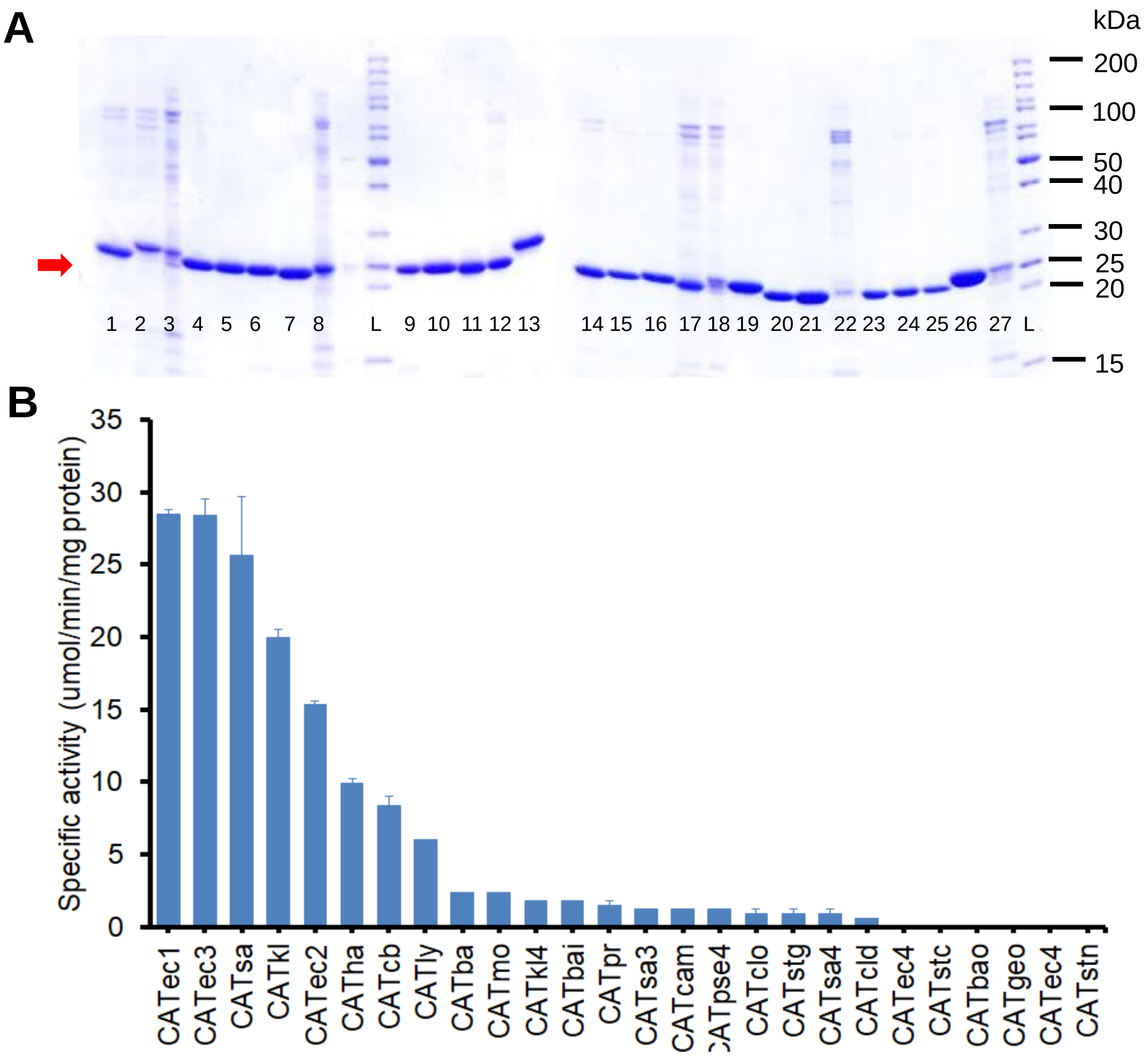

**Figure S3.** Thermostability of the select CATs. Residual activity after the heat incubation at 50ºC was used for the normalization. Each value represents mean ± 1 stdev from three biological replicates.

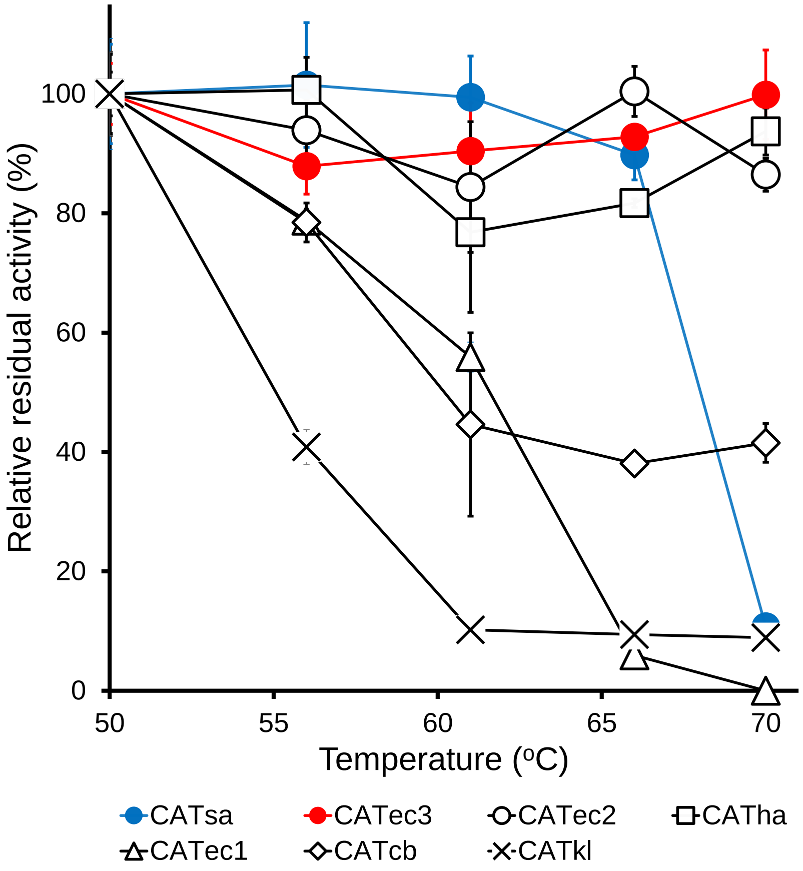

**Figure S4.** Analysis of contacts between atoms of CATsa and CATec3. **(A-B)** Visualized contacts in **(A)** CATsa and **(B)** CATec3. Six types of contacts including hydrogen bonds (H), metal (M), ionic (IH), arene (A), covalent (C), and Van der Waals distance (V) were analyzed in the structures. **(C)** Summarized contact types and amino acid residues contributing the interactions. Energy and Dist denote the interaction energy in kcal/mol and the distance between the centroids of the interacting atoms, respectively. BB denotes whether any of the interacting atoms in the entry are backbone (b) or not (-).

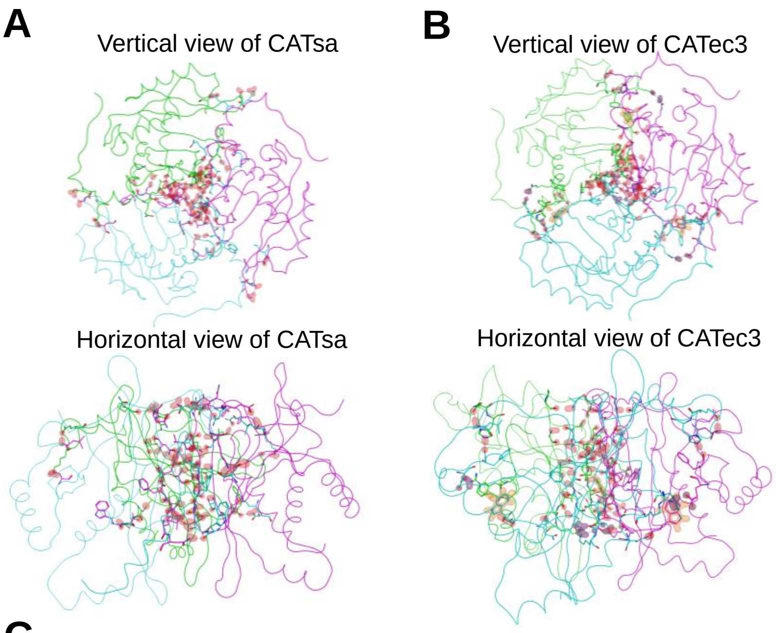

C

CATsa

| Type | Chain A | Chain B | Energy | Dist | BB |
|---|---|---|---|---|---|
| H | Ser94 | Lys12 | -3.6 | 3.13 | -- |
| H | Leu128 | Glu15 | -4.5 | 2.78 | b- |
| H | Trp146 | Asn202 | -2.1 | 3.1 | b- |
| H | Thr147 | Met201 | -1.3 | 3.87 | -- |
| H | Thr150 | Thr30 | -4.5 | 2.83 | bb |
| H | Thr150 | Thr150 | -2.5 | 2.68 | -b |
| H | Gly151 | Thr30 | -2.6 | 3.35 | bb |
| H | Phe152 | Ser28 | -0.6 | 3.37 | bb |
| H | Asn153 | Ser28 | -7.8 | 3 | bb |
| H | Asn155 | Thr26 | -7 | 2.8 | bb |
| H | Asn155 | Asn153 | -6.6 | 3.08 | -- |
| H | Asn157 | Gln23 | -0.5 | 3.49 | -b |
| H | Asn157 | Asn159 | -1.1 | 3.29 | -- |
| | | Total Energy | -44.7 | | |

CATec3

| Type | Chain A | Chain B | Energy | Dist | BB |
|---|---|---|---|---|---|
| IH | Glu95 | Arg199 | -9.42 | 3.03 | -- |
| H | Phe97 | His16 | -5.5 | 2.81 | b- |
| H | Leu128 | Glu15 | -5 | 2.77 | b- |
| A | Trp146 | Phe195 | -1.1 | 4.04 | -- |
| H | Trp146 | Asn202 | -2.4 | 3.06 | b- |
| H | Asn148 | Ser32 | -0.5 | 3.17 | b- |
| H | Asp150 | Thr31 | -4.6 | 2.83 | bb |
| IH | Asp150 | Lys33 | -8.76 | 2.73 | -- |
| H | Ser151 | Thr31 | -0.6 | 3.43 | bb |
| H | Phe152 | Ser29 | -0.7 | 3.36 | bb |
| H | Asn153 | Ser29 | -7.4 | 3.02 | bb |
| H | Asn153 | Asn153 | -0.6 | 3.27 | -- |
| H | Asn155 | Gly27 | -6.9 | 2.86 | bb |
| H | Asn155 | Asn153 | -7 | 3 | -- |
| H | Ala157 | Pro25 | -3.4 | 3.18 | bb |
| | | Total Energy | -63.88 | | |

**Figure S5.** Binding affinity analysis of CATEc3 wildtype and Y20F mutant towards various alcohols. **(A-B)** Docking simulation of various alcohols into the binding pocket of **(A)** CATEc3 WT and **(B)** CATEc3 Y20F. **(C)** Alcohol substrate affinity changes in CATEc3 Y20F compared to CATEc3 WT. △△Affinity (kcal/mol) values were calculated by subtracting △Affinity of CATEc3 WT from △Affinity of CATEc3 Y20F. **(D)** Correlation between the △Affinity and the catalytic efficiency ($k_{cat}/K_M$) of CATEc3 Y20F towards various alcohol substrates.

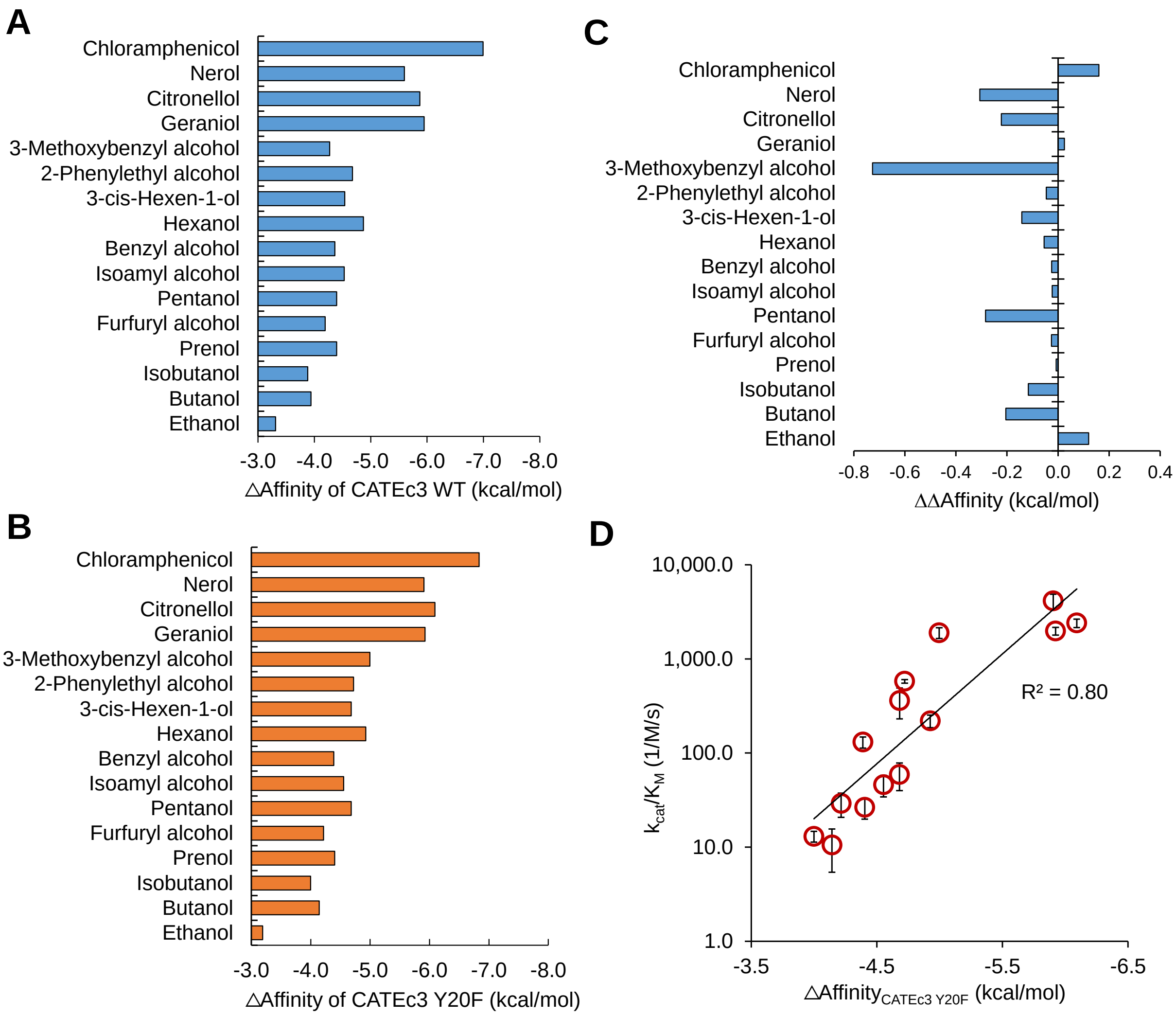

**Figure S6.** *In vitro* fatty acetate ester synthesis by CATec3 Y20F. **(A-C)** Michaelis-Menten plots of CATec3 Y20F towards **(A)** ethanol, **(B)** octanol, and **(C)** decanol. Each data represents mean ± 1 stdev from at least three biological replicates. **(D)** GC/MS chromatograph of acetyltransferase reaction by CATec3 Y20F *in vitro*. The control is a reaction by crude extract of *E. coli* harboring an empty plasmid. The black and red arrows indicate internal standard (n-decane) and decyl acetate (DcAc), respectively. **(E)** Mass fragmentation (m/z) of n-decane and decyl acetate.

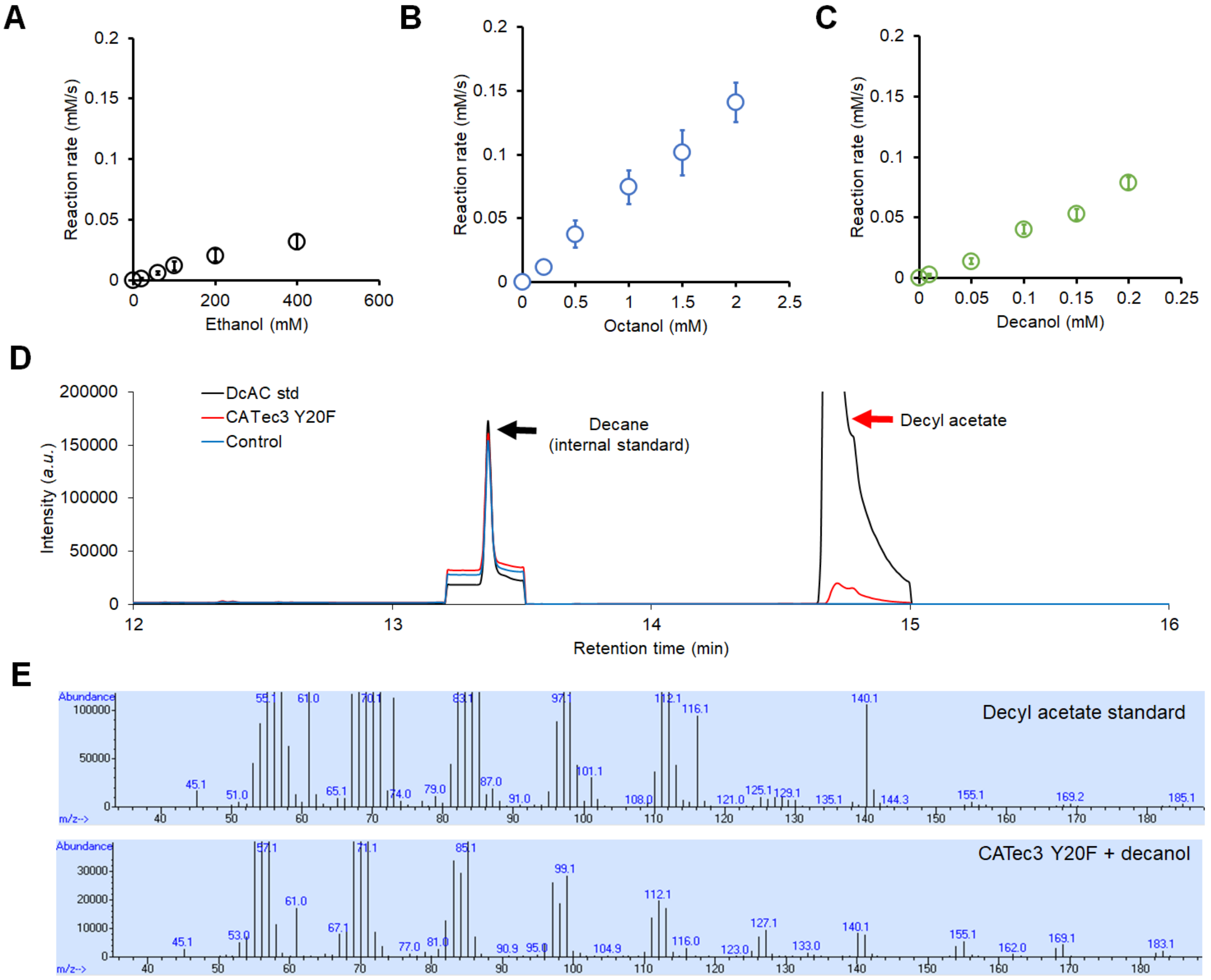

**Figure S7.** *In vitro* specific activity of CATec3 Y20F and CATsa Y20F A138T towards various alcohols. The specific activity was measured with 20mM of alcohol substrates. Each value represents mean ± 1 stdev from three biological replicate.

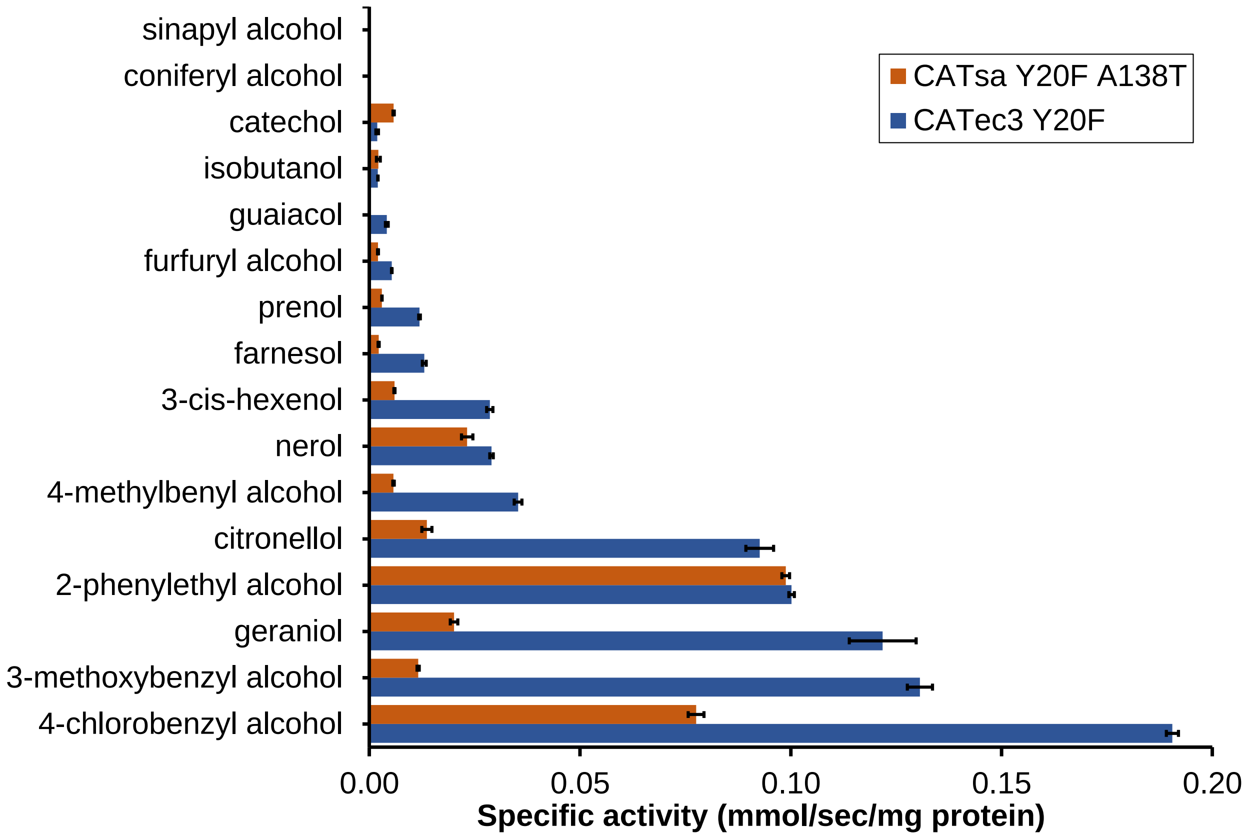

**Figure S8.** Microbial biosynthesis of isoamyl lactate by co-feeding the recombinant *E. coli* co-expressing CATec3 Y20F and propionate CoA transferases (PCTs) with isoamyl alcohol and lactate. **(A)** Scheme of isoamyl lactate microbial biosynthesis. **(B)** Effect of expressing various PCTs on isoamyl lactate production after 48 h. The initial M9 medium contained 10 g/L glucose, 5 g/L yeast extract, 2 g/L of isoamyl alcohol, 2 g/L lactate, and 0.1mM of IPTG to induce the protein expression. 1 mL of hexadecane was overlaid to extract the isoamyl lactate produced during the fermentation. Each value represents mean ± 1 stdev from three biological replicates. Origin of PCTs: PCTpt, *Pelotomaculum thermopropionicum*; PCTme, *Megasphera elsdenii*; PCTtt, *Thermus thermophilus*, PCTcp, *Clostridium propionicum*; PCTre, *Ralstonia eutropha*. **(C)** GC/MS chromatographs of isoamyl lactate microbial biosynthesis. **(D)** Mass fragmentation of isoamyl lactate.

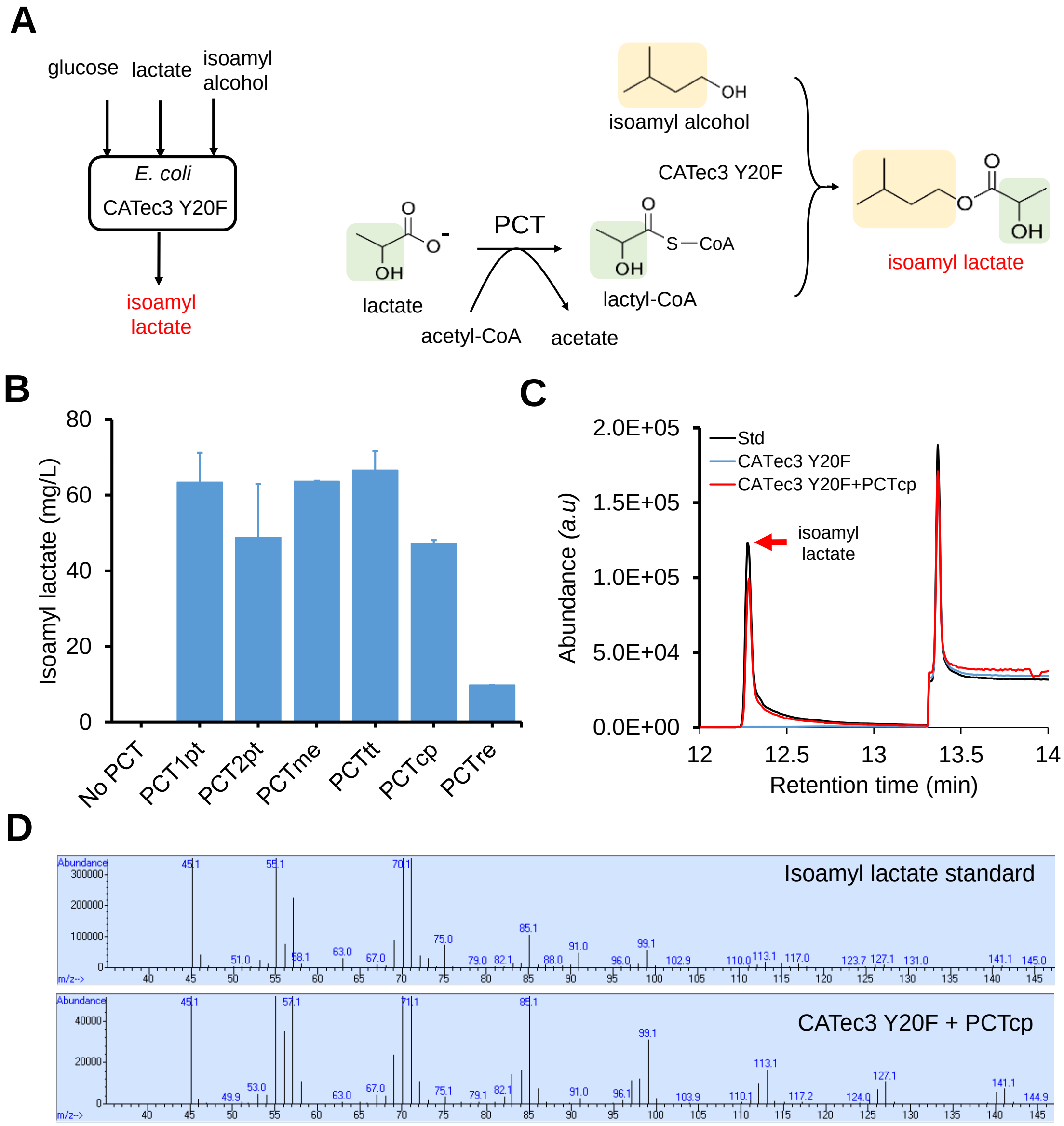

**Figure S9.** Production of acetate esters by co-feeding the recombinant *E. coli* harboring CATec3 Y20F with glucose and alcohols at 37$^{o}$C. The alcohols were supplemented to the initial M9 glucose medium at concentrations of 3 g/L for butanol, isobutanol, pentanol, and isoamyl alcohol, 1 g/L for benzyl alcohol, and phenylethyl alcohol, and 0.3 g/L for geraniol. Each value represents mean ± 1 stdev from three biological replicates.

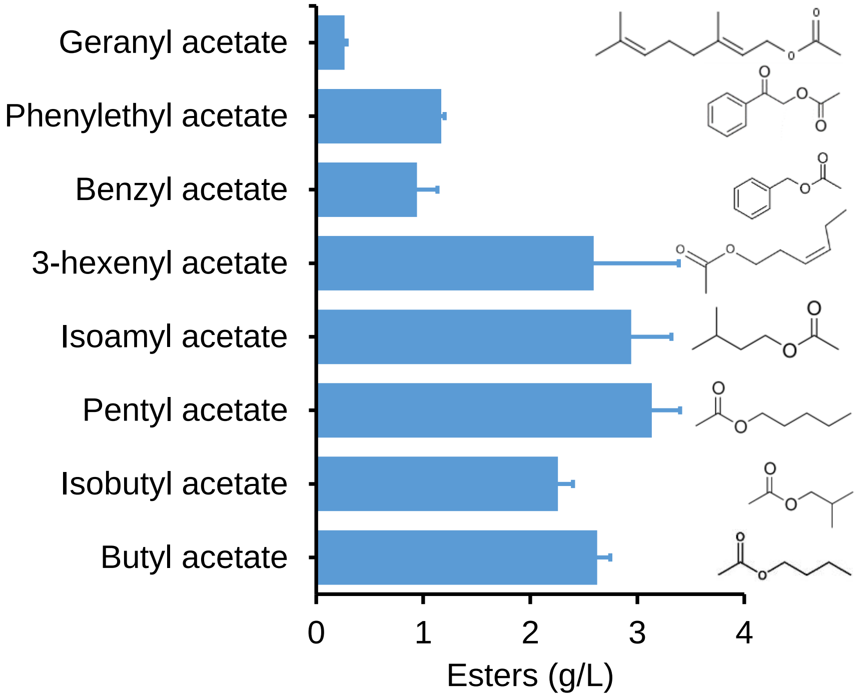

**Figure S10:** Fed-batch fermentation coupled with *in situ* ester extraction for conversion of **(A)** isoamyl alcohol to isoamyl acetate and **(B)** 2-phenylethyl alcohol to 2-phenylethyl acetate. Each value represent mean ± 1 stdev from three biological replicates.

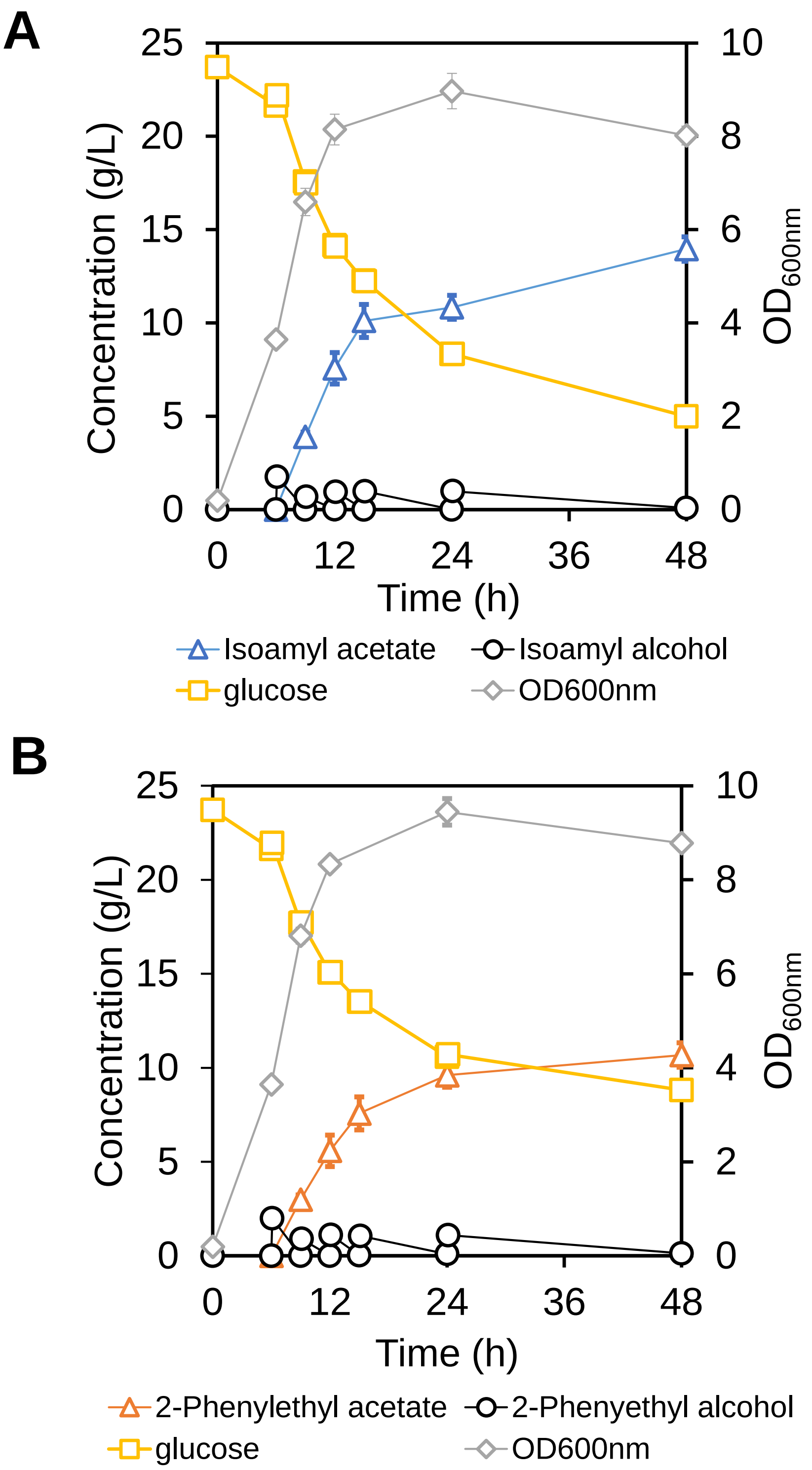

**Figure S11.** Microbial biosynthesis of an ester profile from roses by a recombinant CATec3 Y20F-expressing *E. coli* HSEC01. **(A)** Conversion of an alcohol mixture to an ester profile with hexadecane overlay during fermentation. **(B)** Cell growth kinetics during the alcohol mixture conversion. The arrow indicates the point when the alcohol mixture was added to the medium. Each value represents mean ± 1 stdev from three biological replicates.

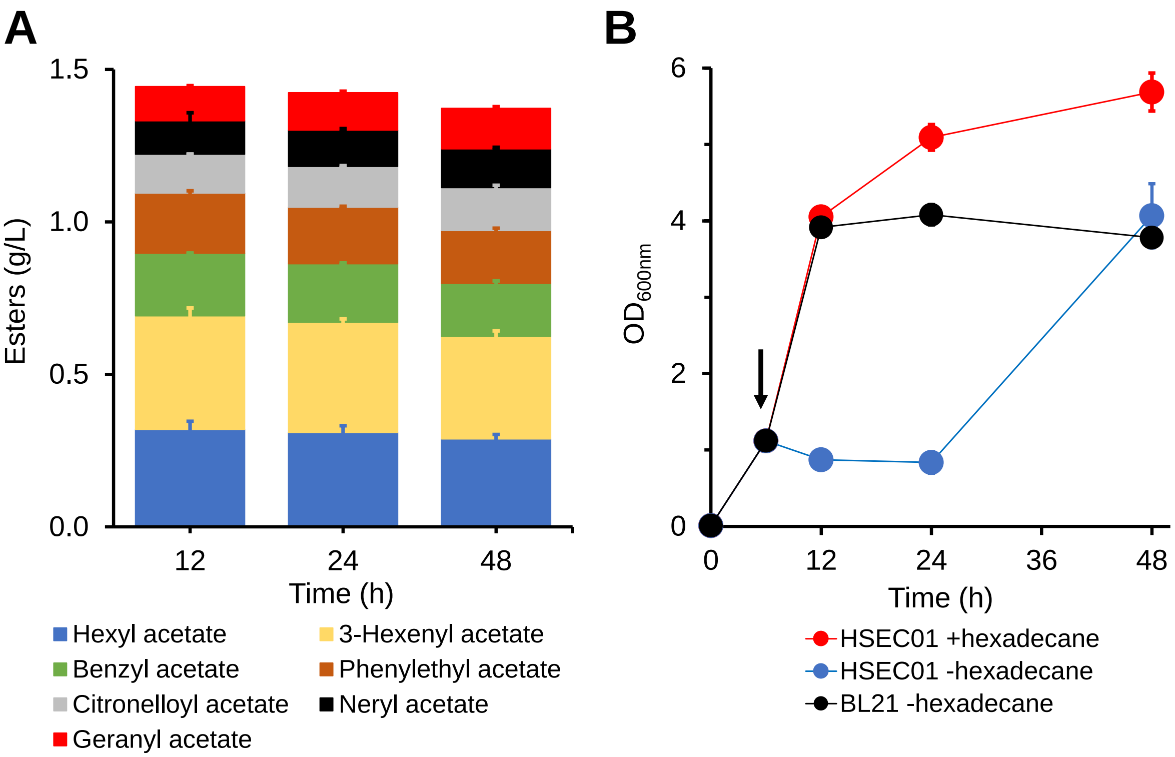

**Figure S12.** Production of acetate esters by co-feeding the recombinant *C. thermocellum* harboring CATec3 Y20F with glucose and alcohols at 55°C. Kinetic profiles of biosynthesis of **(A)** designer bioesters and **(B)** isobutyrate esters as byproducts. The arrow indicates the time point when the alcohol mixture was added to the medium. Each value represents mean ± 1 stdev from three biological replicates.

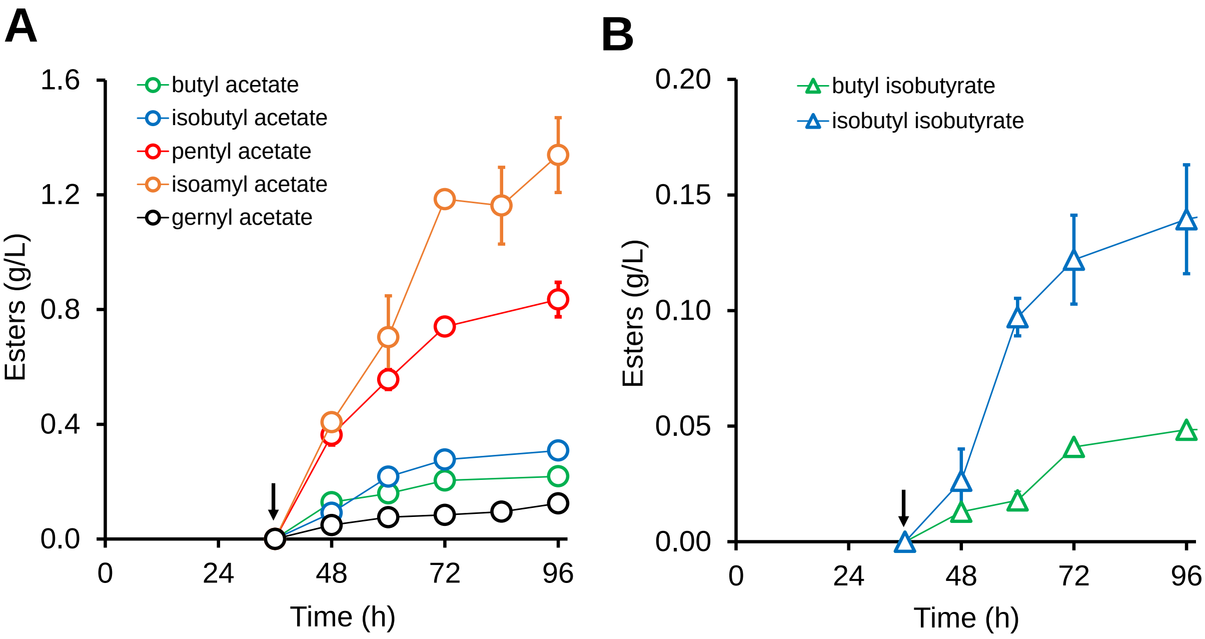

**Figure S13.** Toxicity effect of isobutanol on microbial health of *C. thermocellum*. The *C. thermocellum* DSM1313 Δ*hpt* strain (M1354) was cultured in MTC medium containing 5g/L cellobiose and isobutanol. Each value represents mean ± 1 stdev from three biological replicates.

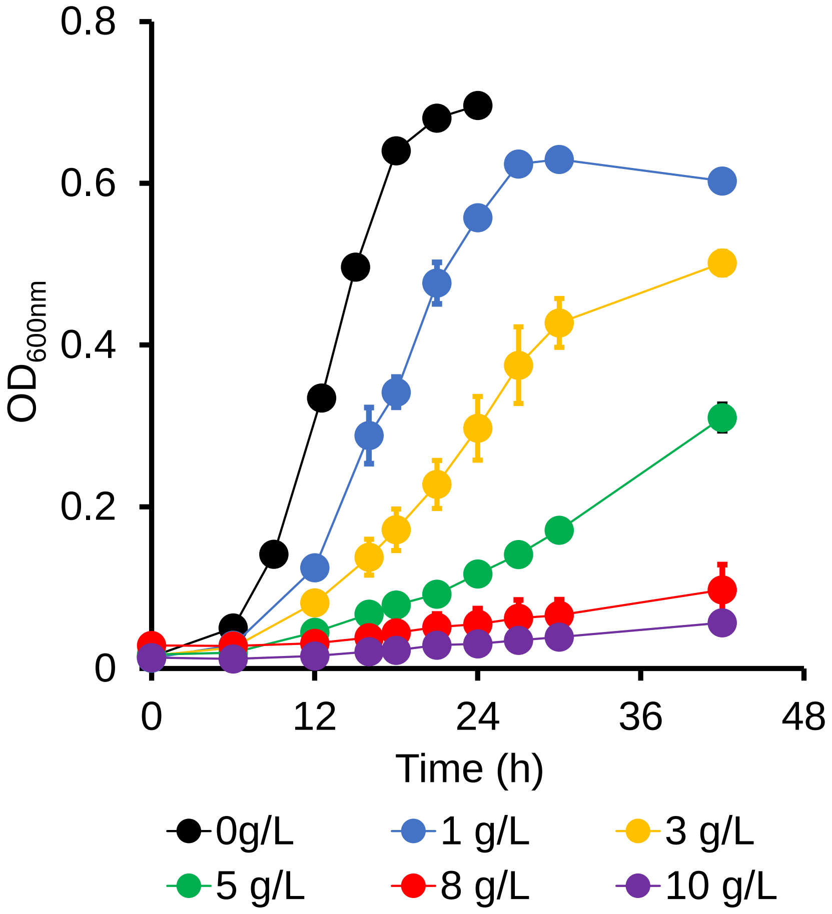

**Figure S14.** Proteome analysis of CATec3 and CATec3 Y20F expressing *C. thermocellum* **(A)** Peptide fragmentations of CATec3 used for the proteomics analysis. **(B)** Relative proteome abundance of HSCT2108 (CATec3 Y20F) versus HSCT2107 (CATec3). The red dot indicates the increased abundance of CATec3 Y20F in HSCT2108. Each value represents the significance (-$\log_{10}$ p-value; y-axis) and average difference (in $\log_2$; x-axis) between HSCT2108 and HSCT2017 across three biological replicates.

**A**

| Peptide fragments |
|---|
| LLLPLSVQVHHAVCDGFHVAR.1xCarbamidomethyl [C14] |
| LPCGFSLTSK.1xCarbamidomethyl [C3] |
| EHFEFXR |
| LQELCNSK.1xCarbamidomethyl [C5] |
| SLDDSAYK |
| IDITTLK |

**B**

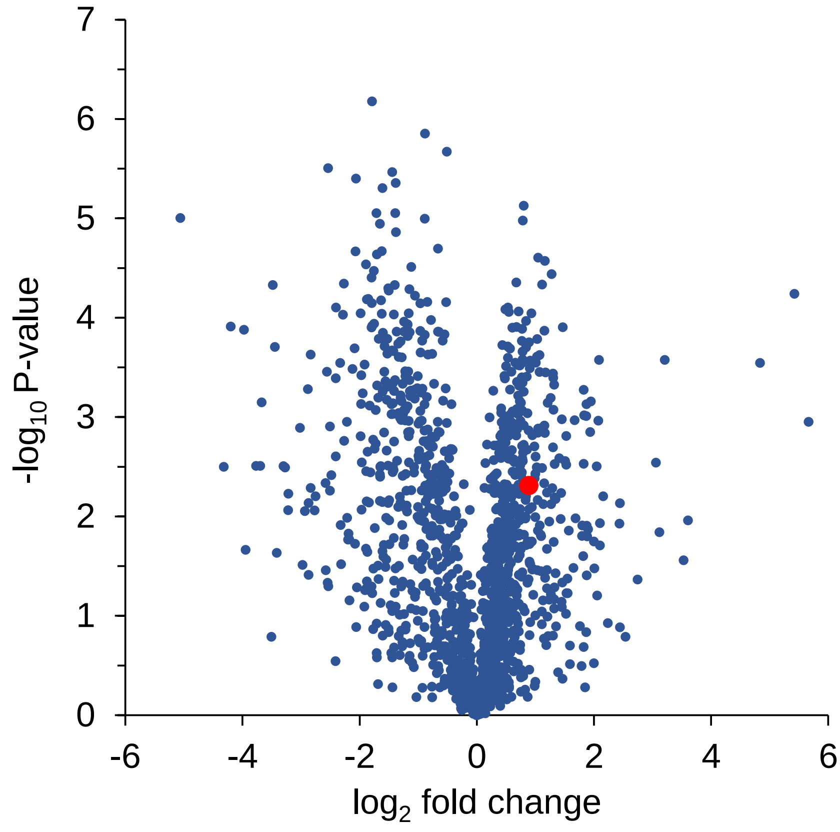
